# Resistance Gene Association and Inference Network (ReGAIN): A Bioinformatics Pipeline for Assessing Probabilistic Co-Occurrence Between Resistance Genes in Bacterial Pathogens

**DOI:** 10.1101/2024.02.26.582197

**Authors:** Elijah R. Bring Horvath, Mathew G. Stein, Matthew A. Mulvey, Edgar J. Hernandez, Jaclyn M. Winter

**Affiliations:** Department of Pharmacology and Toxicology, University of Utah, Salt Lake City, Utah, 84112, United States; Department of Medicinal Chemistry, University of Utah, Salt Lake City, Utah, 84112, United States; School of Biological Sciences, University of Utah, Salt Lake City, UT 84112, United States; Henry Eyring Center for Cell & Genome Science, University of Utah, Salt Lake City, UT 84112, United States; Department of Biomedical Informatics, University of Utah, Salt Lake City, Utah, 84112, United States

## Abstract

The rampant rise of multidrug resistant (MDR) bacterial pathogens poses a severe health threat, necessitating innovative tools to unravel the complex genetic underpinnings of antimicrobial resistance. Despite significant strides in developing genomic tools for detecting resistance genes, a gap remains in analyzing organism-specific patterns of resistance gene co-occurrence. Addressing this deficiency, we developed the Resistance Gene Association and Inference Network (ReGAIN), a novel web-based and command line genomic platform that uses Bayesian network structure learning to identify and map resistance gene networks in bacterial pathogens. ReGAIN not only detects resistance genes using well- established methods, but also elucidates their complex interplay, critical for understanding MDR phenotypes. Focusing on ESKAPE pathogens, ReGAIN yielded a queryable database for investigating resistance gene co-occurrence, enriching resistome analyses, and providing new insights into the dynamics of antimicrobial resistance. Furthermore, the versatility of ReGAIN extends beyond antibiotic resistance genes to include assessment of co-occurrence patterns among heavy metal resistance and virulence determinants, providing a comprehensive overview of key gene relationships impacting both disease progression and treatment outcomes.

**Graphical Abstract:** 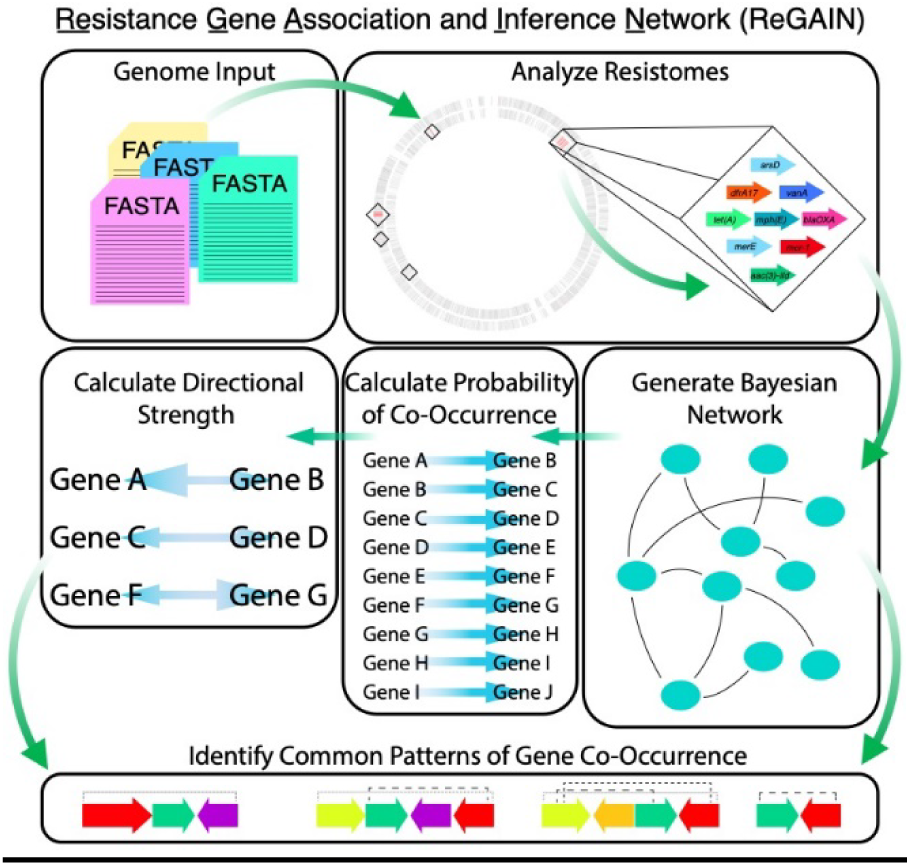

## Introduction

Increasing rates of multidrug resistance in clinically important bacterial pathogens represents a monumental threat to global human health (1–4). Among antibiotic resistant (AR) bacteria, those classified as multidrug resistant (MDR) represent the most challenging health threat (5,6), as they can encode extensive genetic machinery to evade elimination by anti-infective agents (7–9), thereby significantly limiting treatment options. Continued increases in rates of MDR bacteria has been historically attributed to improper management of antibiotics, including misuse of antibiotics in treating non-bacterial infections and antibiotic overuse in agriculture (10–13). Moreover, the increase in multidrug resistance is significantly driven by the dissemination and acquisition of resistance genes within microbial communities through horizontal gene transfer (14–21). Indeed, many multidrug resistance gene operons are flanked by transposable elements, further indicating the mobility of not just small operons, but large genomic islands containing extensive resistance gene diversity (7,22–25). The ability of bacteria to easily exchange antibiotic resistance genes (ARGs) makes antibiotic and multidrug resistance an ever-evolving problem (10,26). Furthermore, exposure to heavy metals in the environment, combined with genetically encoded heavy metal resistance genes (HMGs), could aid in the transfer of ARGs through co-selection (27–29). To better monitor the spread and dissemination of resistance genes, careful curation of genomic datasets, including isolation source, region, and date should be a priority.

With advancements in whole-genome sequencing, a wealth of bacterial genomic data has become available, offering unprecedented opportunities to understand the evolution of multidrug resistance and map common patterns of resistance gene co-occurrence in problematic bacterial species. This is especially important in monitoring patterns of co-occurrence with genes conferring resistance to last-line antibiotics, such as the *mcr*-family of colistin resistance genes (16,30,31). Though powerful tools have been developed to identify antibiotic resistance genes, e.g., AMRfinderPlus and ResFinder (32,33), these tools are predominantly confined to gene identification, overlooking the complex interplay between genes that contribute to the MDR phenotype. Several *in silico* methods designed to expand the analysis of resistome data have been published. However, these methods generally require researchers to have extensive computational experience (27,34,35) or focus on resistance gene abundance within metagenomic datasets rather than organism-specific patterns of resistance gene co-occurrence (35,36). Currently, to the best of our knowledge, there is not a publicly available bioinformatic platform designed to measure antibiotic resistance gene co-occurrence.

In response to the urgent need to not only identify resistance genes but also catalog patterns of resistance gene co-occurrence, we developed the <u>Re</u>sistance <u>G</u>ene <u>A</u>ssociation and Inference <u>N</u>etwork (ReGAIN) genomic pipeline (Figure 1). Leveraging the foundational capabilities of the National Center for Biotechnology Information (NCBI) software AMRfinderPlus (32), ReGAIN’s core pipeline employs a robust Bayesian network structure learning approach to elucidate probabilistic patterns of resistance or virulence gene co-occurrence indicative of multidrug resistance in microbes. The ReGAIN pipeline is not limited to resistance genes and extends to virulence determinants through a parallel pathway, thus providing a comprehensive view of microbial defense mechanisms. Designed to be flexible and user-friendly, the ReGAIN web application simplifies the bioinformatic workflow into two core modules: data acquisition and Bayesian network analysis. Users are directed to upload genome files in FASTA format to the data acquisition module, specify an analytical focus (resistance or virulence), and initiate the analysis through a simple submission form. Using externally prepared data or files generated from the data acquisition module, users can then upload data to the Bayesian network analysis module. This module provides interactive Bayesian networks and probabilistic measurements, like conditional probability and relative risk ratios, along with confidence intervals and standard deviation. Further distinguishing the ReGAIN pipeline are two novel post-hoc analyses that compute the bidirectional strength of gene co-occurrence, which offer a quantitative method to explore and interpret asymmetry in gene-gene relationships. These metrics enhance our understanding of resistance gene networks, offering new insights into the dynamics of multidrug resistance. For researchers who prefer to have more control over the analyses, ReGAIN is additionally being developed as a command line program.

**Figure 1.**
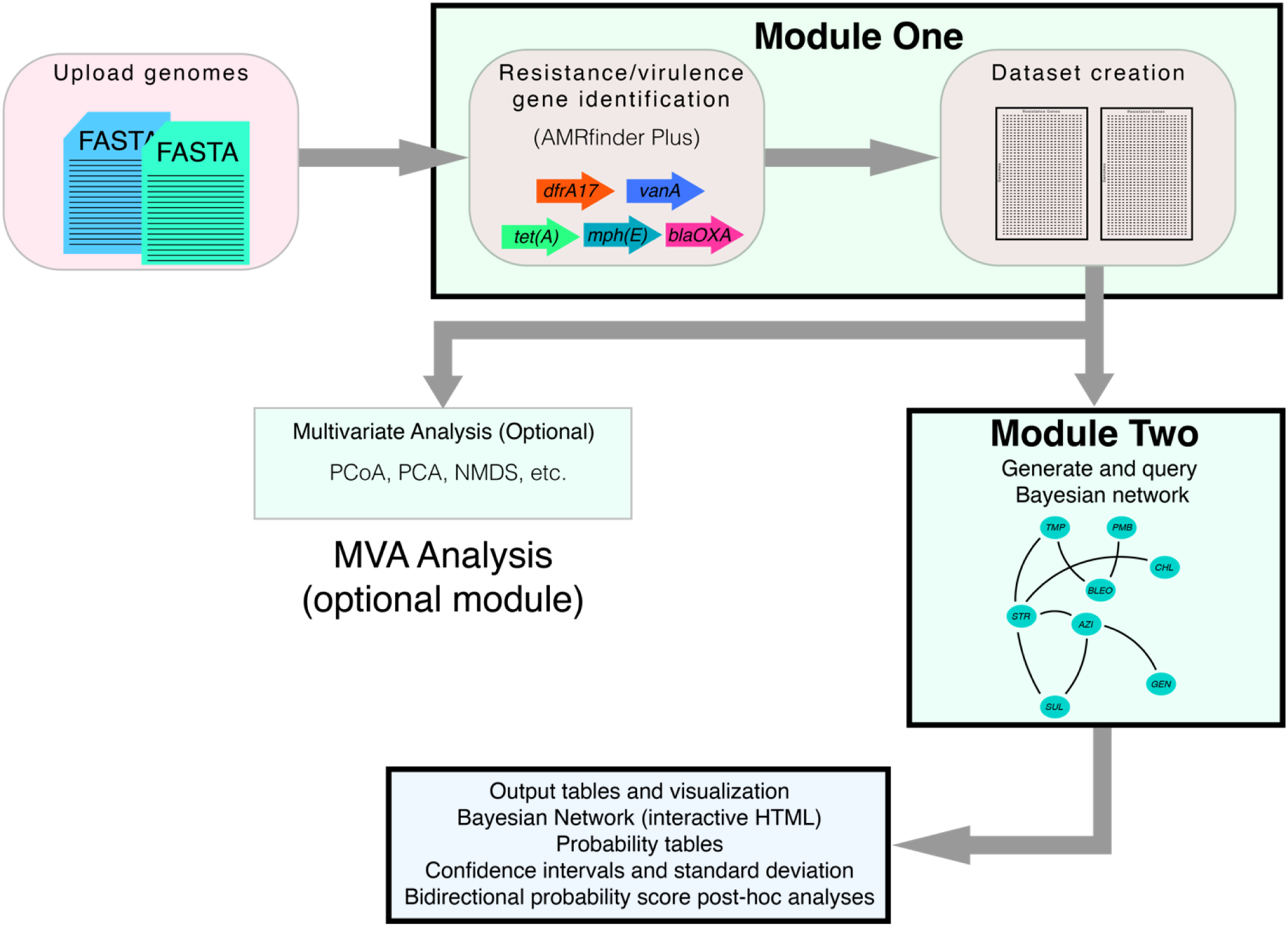
Outline of the ReGAIN bioinformatic pipeline. The first core module uses AMRfinder Plus to identify resistance/virulence genes from the input genomes in nucleotide FASTA format. After all genomes have been processed, two major output CSV files are created: a presence/absence matrix of genes across genomes and a metadata file containing all identified genes and their associated resistance type. All downstream ReGAIN analyses can be performed using these two output files. The second core module, the Bayesian analysis pipeline, utilizes 100 resamples of the submitted dataset to generate a table of pairwise conditional probability and relative risk scores for each gene pair in the dataset, including low and high confidence intervals and standard deviation. Additional post-hoc analyses generate bidirectional probability scores based on both conditional probability and relative risk for each gene pair cohort. Both tables are output in CSV format. Finally, an interactive Bayesian network HTML web page is generated. The optional additional analyses module includes Association Rule Mining and multivariate analyses.

As an initial step towards identifying patterns of resistance gene co-occurrence, Bayesian networks were constructed using publicly available genomic data for a diverse set of common, clinically important bacterial pathogens. These included *Escherichia coli* and *Enterococcus faecalis* and the ESKAPE pathogens *Enterococcus faecium, Staphylococcus aureus*, *Klebsiella pneumoniae*, *Acinetobacter baumannii*, *Pseudomonas aeruginosa*, and *Enterobacter* spp., which are responsible for the majority of nosocomial infections worldwide and are often resistant to multiple antibiotics(1,6,8,13,37–39). Our large-scale Bayesian network analyses offer valuable insight into general patterns and pairwise probabilities of resistance gene co-occurrence among pathogens and provides a platform that can also reveal patterns of heavy metal resistance and virulence gene co-occurrence.

## Results

### Overview of ReGAIN

The ReGAIN platform is comprised of two primary modules: a data acquisition and dataset construction module and a Bayesian network structure learning module. The front-end input of the ReGAIN web server data acquisition module allows users to upload assembled bacterial genomes in FASTA format. After genome submission, users are directed to select an organism-specific or non-specific pipeline. The organism-specific pipeline takes advantage of the AMRfinderPlus software (32), which analyzes the genomic dataset for organism-specific resistance-conferring point mutations and genes. Currently, the ReGAIN organism-specific pipeline supports *Acinetobacter baumannii, Burkholderia cepacian, Burkholderia pseudomallei, Camplylobacter* spp.*, Clostridium difficile, Enterococcus faecalis, Enterococcus faecium, Escherichia* spp., *Klebsiella* spp., *Neisseria* spp., *Pseudomonas aeruginosa, Salmonella* spp., *Staphylococcus aureus, Staphylococcus pseudintermedius, Streptococcus agalactiae, Streptococcus pneumoniae, Streptococcus pyogenes,* and *Vibrio cholerae* (Table 1). Additional options allow users to set a minimum/maximum gene occurrence threshold, as the exclusion of very low or high abundance genes is an important step in reducing network noise. Because discrete Bayesian network analyses require variables to exist in at least two states within a dataset, it is necessary to exclude ubiquitously occurring genes from the analysis. The ReGAIN data acquisition workflow creates a presence/absence matrix of genes across the genomic population (‘1’ = present, ‘0’ = absent). Thus, for ReGAIN datasets, any given gene must occur in both the ‘present’ and ‘absent’ states. To exclude ubiquitously occurring genes while still allowing user-end flexibility, ReGAIN permits users to set the minimum/maximum values themselves; for *n* genomes, the maximum value for any given gene must be at least *n* − 1. From here, users may choose to assess their genomes for resistance or virulence genes. Resistance genes include antibiotic and heavy metal resistance genes, as well as multiple stress response genes.

**Table 1.** Genomic sample size of ReGAIN databases. Database size describes the number of genomes and number of antibiotic and resistance genes included in the database.

| Organism | Database Size |
| --- | --- |
| *Acinetobacter baumannii* | 747 genomes; 105 resistance genes |
| *Clostridium difficile* | 275 genomes; 31 resistance genes |
| *Enterobacter cloacae* | 623 genomes; 62 resistance genes |
| *Enterococcus faecalis* | 1064 genomes; 58 resistance genes |
| *Enterococcus faecium* | 502 genomes; 39 resistance genes |
| *Escherichia coli* | 1491 genomes; 179 resistance genes |
| *Klebsiella pneumoniae* | 2464 genomes, 142 resistance genes |
| *Neisseria gonorrhoeae* | 173 genomes; 43 resistance genes |
| *Pseudomonas aeruginosa* | 867 genomes; 158 resistance genes |
| *Salmonella enterica* | 2391 genomes; 118 resistance genes |
| *Salmonella enterica* subsp. *typhi* | 158 genomes; 16 resistance genes |
| *Staphylococcus aureus* | 1678 genomes; 91 resistance genes |
| *Streptococcus pneumoniae* | 390 genomes; 11 resistance genes |

Using the NCBI AMRfinderPlus software, user-uploaded genomic datasets can be assessed for the presence of core antibiotic resistance genes (organism non-specific analysis) or for core resistance genes along with species-specific point mutations and virulence genes (organism specific analysis). AMRfinderPlus utilizes both Hidden Markov Models and manually curated BLAST cutoffs to identify resistance and virulence genes based on the large Reference Gene Catalog consisting of over 6,000 genes (32). Identification of genes and creation of data files required for downstream analysis is automated within the data acquisition workflow. First, the distribution of genes across the genomic dataset is reduced to a binary format and a presence/absence data matrix is built. This data matrix is then curated based on the user-input minimum/maximum gene occurrence values. Finally, a metadata file is generated, which contains a list of all genes identified in the analysis, independent of minimum/maximum gene values. For resistance genes, additional information associated with gene function, such as resistance type or gene class (i.e., aminoglycoside, sulfonamide, β-lactamase, etc.) will be included. Output files from the data acquisition module include results files for each uploaded genome, an initial data matrix, a curated final data matrix, and a metadata file. From here, the final data matrix and metadata file can be submitted to the Bayesian network structure learning module.

### Bayesian Network Structure Learning Analysis

The Bayesian network analysis module has been optimized to handle both very large datasets (≥ 100 genes) and small-to-moderate sized datasets (<100 genes). The module’s backend employs the bnlearn (40) and gRain (41) package in R. Given the computational complexity associated with Bayesian network queries, two pipelines were designed based on the size of the input dataset to optimize performance and accuracy. For small-to-moderate size datasets, ReGAIN utilizes the gRain ‘queryGrain’ function. This function is adept at handling smaller networks where exhaustive computations can be performed without prohibitive computational costs and full graph inference is manageable, thus allowing direct computation of conditional dependencies. Conversely, for very large datasets, we leverage ‘cpquery’, a function designed for large-scale Bayesian networks. As variables increase, the computational requirements for direct calculations increase exponentially, quickly resulting in intractable computation costs. ‘cpquery’ addresses this issue by estimating conditional probabilities through Monte Carlo simulations that scale linearly with the number of variables, thereby facilitating the analysis of large genomic networks (40,42,43). Altogether, these methods ensure that our pipeline remains computationally feasible while maintaining a high degree of accuracy in the probabilistic assessment of gene co-occurrence. Although the Bayesian network analysis module is designed to interface with the results output from the data acquisition module, instructions on how to format externally prepared datasets are provided for users to run this module independently of the Data Acquisition pipeline (Figure S1).

Once executed, the Bayesian network analysis module employs a systematic approach to analyze the genomic data. First, Bayesian network structure learning using a Hill Climbing algorithm and Bayesian Dirichlet equivalent score is used to construct the network. The combination of these two functions guides the algorithm in learning the most probable network structure given the data. A critical aspect of the Bayesian network workflow is the implementation of bootstrapping to resample data multiple times. Users are directed to use between 300 and 500 bootstraps to enhance the robustness of the network analysis. Following bootstrapping, the module applies a significance threshold of 0.5 to filter out weakly supported gene pairs from the network. The refined network then undergoes further analysis through a resampling process. This step involves fitting multiple Bayesian networks to subsets of the data, which are randomly sampled with replacement, such that all values within the dataset have an equal probability of being selected one or multiple times. To enhance the statistical reliability of the results, the resampled datasets were used 100 times to fit the Bayesian network. Finally, statistical analyses are performed on the fitted networks. For each gene pair, median conditional probability (Equation 1), median relative risk (Equation 2), empirical confidence intervals, and standard deviation are calculated. Conditional probability is described as the probability of observing Gene A given the presence of Gene B, *P(A|B)*, while relative risk is the ratio of the conditional probability of observing Gene A given Gene B to the conditional probability of only observing Gene A in the absence of gene B, *P(A|B)/P(A|*­*B)*. Whereas conditional probability is expressed on a scale of 0 to 1 (e.g., a conditional probability of 0.65 indicates a 65% probability of observing the gene pair), relative risk offers more insight. As relative risk is a ratio, it offers a more descriptive scale. For instance, a relative risk of 1 suggests that Gene A and Gene B are likely independent of each other, A **⫫** B. A value greater than 1 suggests that it is more likely to observe Gene A in the presence of Gene B, as *P(A|B) > P(A|*­*B).* Similarly, a value less than 1 indicates that Gene A is more likely to be observed in the absence of Gene B, as *P(A|B) < P(A|*­*B)*. Aside from a table containing statistical summaries, conditional probability, and relative risk values, the output of the Bayesian network analysis module includes the refined network as an interactive HTML file.

To further analyze the probabilistic gene co-occurrence relationships, two post-hoc analyses are performed. Bidirectional probability score (BDPS) (Equation 3) describes the directional strength of the conditional probabilities of a gene pair. As conditional probability, *P(A|B)*, is dependent on the presence or absence of Gene B, the reciprocal conditional probability is not often equal, *P(A|B) ≠ P(B|A)*. Therefore, a better assessment of the overall strength of the relationship can be achieved by taking the ratio of these two values. If BDPS = 1, equal bidirectional strength can be assumed. If BDPS > 1, the probability of observing Gene A given Gene B is stronger, *P(A|B) > P(B|A)*. Conversely, if BDPS < 1, the probability of observing Gene B given Gene A is stronger, *P(A|B) < P(B|A)*. Using relative risk, fold change can be calculated (Equation 4), an additional post-hoc analysis interpreted similarly to BDPS, and these scores are output to a second CSV file. Fold change is interpreted inversely to BDPS; due to the nature of how relative risk ratios are calculated, a fold change > 1 indicates that the probability of observing Gene A given Gene B is weaker, RR(A|B) < RR(B|A), while a fold change < 1 indicates that the probability of observing Gene A given the presence of Gene B is stronger, RR(A|B) > RR(B|A). Additionally, a fold change value of 1 may indicate either equal bidirectional probability or variable independence, dependent on the constituent relative risk values. To expand on this, if Gene A and Gene B and the reciprocal both exhibit a relative risk of 1, the fold change would also be 1. Because of this, it is important to use these post-hoc analyses to supplement conditional probability and relative risk of each gene pair. Taken together, the Bayesian network structure learning module offers valuable insight into both the probability and directional strength of gene co-occurrence.

### Additional Statistical Analyses

In addition to Bayesian network structure learning, the ReGAIN platform offers a multivariate analysis module (MVA) for exploring datasets. While Bayesian networks can reveal probabilistic relationships and dependencies between variables, MVA can quickly provide insight into the patterns and structures of variables within the dataset. Further, the use of a k-means clustering algorithm allows for the categorization of data into clusters based on similarity measures, which is independent of the probabilistic models and may assist in identifying distinct groups or outliers. This can be an important analysis in understanding the spread and development of resistance patterns, especially as a dataset grows over time and becomes more complex.

The ReGAIN MVA module begins by constructing a distance matrix dependent on a user-defined distance measurement, including, but not limited to, Jaccard, Bray-Curtis, Euclidean, and Manhattan measures of distance. Principle coordinates analysis (PCoA) is then applied to the distance matrix to reduce dimensionality, facilitating a visual exploration of the data. The percentage variance explained by the first two principal coordinates is calculated to guide interpretation. A k-means clustering algorithm further categorizes the data into user-defined clusters, which incorporate confidence ellipses around clusters for statistical significance to add an additional layer of validation to the analysis (Figure 2) (44). The resulting MVA graphical representation serves as a tool for visual exploration of gene co-occurrence patterns and is designed to complement the Bayesian network structure learning module by providing initial insight into a dataset.

**Figure 2.**
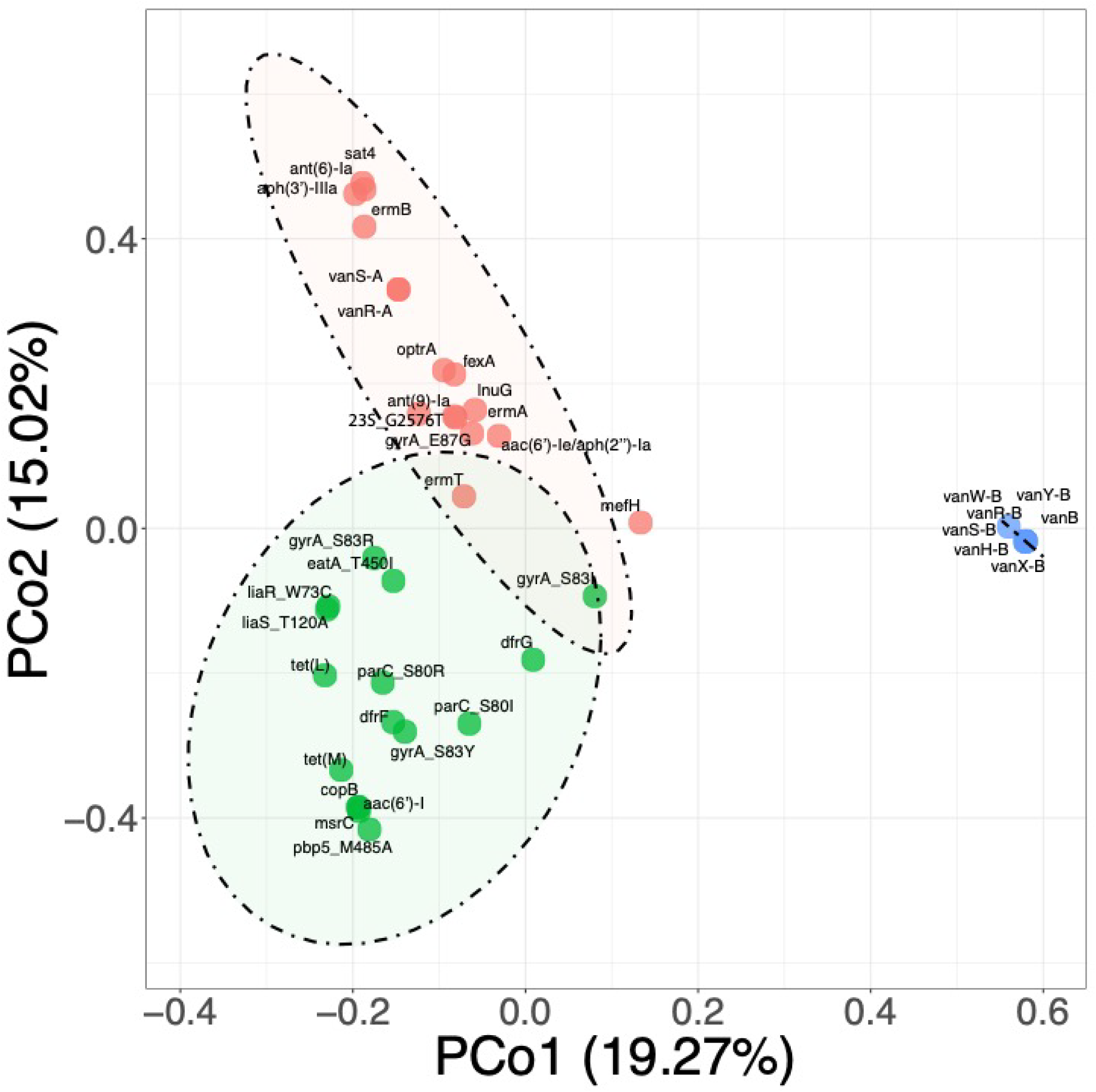
Example Principal Coordinate Analysis (PCoA) output from ReGAIN’s Additional Analyses Module of 39 resistance genes identified in a genomic population of 502 *Enterococcus faecium* genomes. PCoA plot was generated using the Jaccard measure of distance. Ellipses represent 95% confidence, with three clusters specified.

### Probabilistic Resistance Gene Co-Occurrence in Gram-Negative Pathogens

To assess the effectiveness of the the ReGAIN pipeline, we investigated the co-occurrence of resistance genes in thirteen clinically relevant Gram-positive and Gram-negative bacterial species. This analysis involved between 15 and 2464 genomes per species (Figures 3 and 4, Table 1). Initially, each genomic population was examined for the presence of resistance genes (Figure S2, Tables S1 and S2). Bayesian networks were then constructed with a focus on Gram-negative pathogens. Using *E. coli* as a proof of concept, we curated a dataset containing 179 antibiotic and heavy metal resistance genes across 1491 genomes (Figure S2A), resulting in 30802 gene−gene comparisons (Figure 3, Table S1A). To test the accuracy of our system, we analyzed the probabilistic relationship of antibiotic resistance genes that have been previously observed to occur together in Enterobacteriaceae. These genes include the streptomycin resistance genes *aph*(*6*)*-Id* and *aph(3’’)-Ib*, and the sulfonamide resistance gene, *sul2* (9,45,46). In *E. coli*, strong positive probabilistic relationships were observed between *aph(3’’)-Ib* to *aph(6)-Id* (cond. prob. 0.99, rel. risk 89.8), *aph(3’’)-Ib* to *sul2* (cond. prob. 0.99, rel. risk 22.2), and *aph(6)-Id* to *sul2* (cond. prob. 0.99, rel. risk 22.0). Additionally, bidirectional probability scores of 1.02 were observed between *aph(3’’)-Ib* and *sul2*, and 1.01 between *aph(6)-Id* and *sul2*, indicating that the probability of observing either *aph(3’’)-Ib* or *aph(6)-Id*, given the presence of *sul2*, or vice versa, is equally likely. Similar relationships between these genes in *Salmonella enterica* and *K. pneumoniae* were also observed. In *Enterobacter cloacae*, *aph(3’’)-Ib* and *aph(6)-Id* exhibited low conditional probabilities of co-occurrence with *sul2* but displayed strong relative risk scores (cond. prob. 0.27/0.27, rel. risk. 10.9/11.0). This suggests that when adjusted for single-occurrence of either *aph(3’’)-Ib* or *aph(6)-Id*, there is indeed a positive probabilistic relationship between streptomycin and sulfonamide resistance genes in *E. cloacae.* In *A. baumannii*, only *aph(6)-Id* and *sul2* exhibited a positive probabilistic relationship (cond. prob. 0.78, rel. risk 3.0, BDPS 1.3). Conversely, neutral probabilities of co-occurrence were documented between these genes in *P. aeruginosa*, with relative risk scores of 1.02 and 1.05 for *aph(3’’)-Ib* and *sul2* and *aph(6)-Id* and *sul2*, respectively. Interestingly, strong probabilistic relationships were observed between both *aph(3’’)-Ib* and *aph(6)-Id* and the cephalosporin resistance gene, *bla_PER-1_*(cond. prob. 0.96/0.97, rel. risk 19.0/18.9, respectively). These findings indicate that *P. aeruginosa* deviates from the general pattern of co-occurring streptomycin-sulfonamide resistance genes observed in other pathogenic Gammaproteobacteria species.

**Figure 3.**
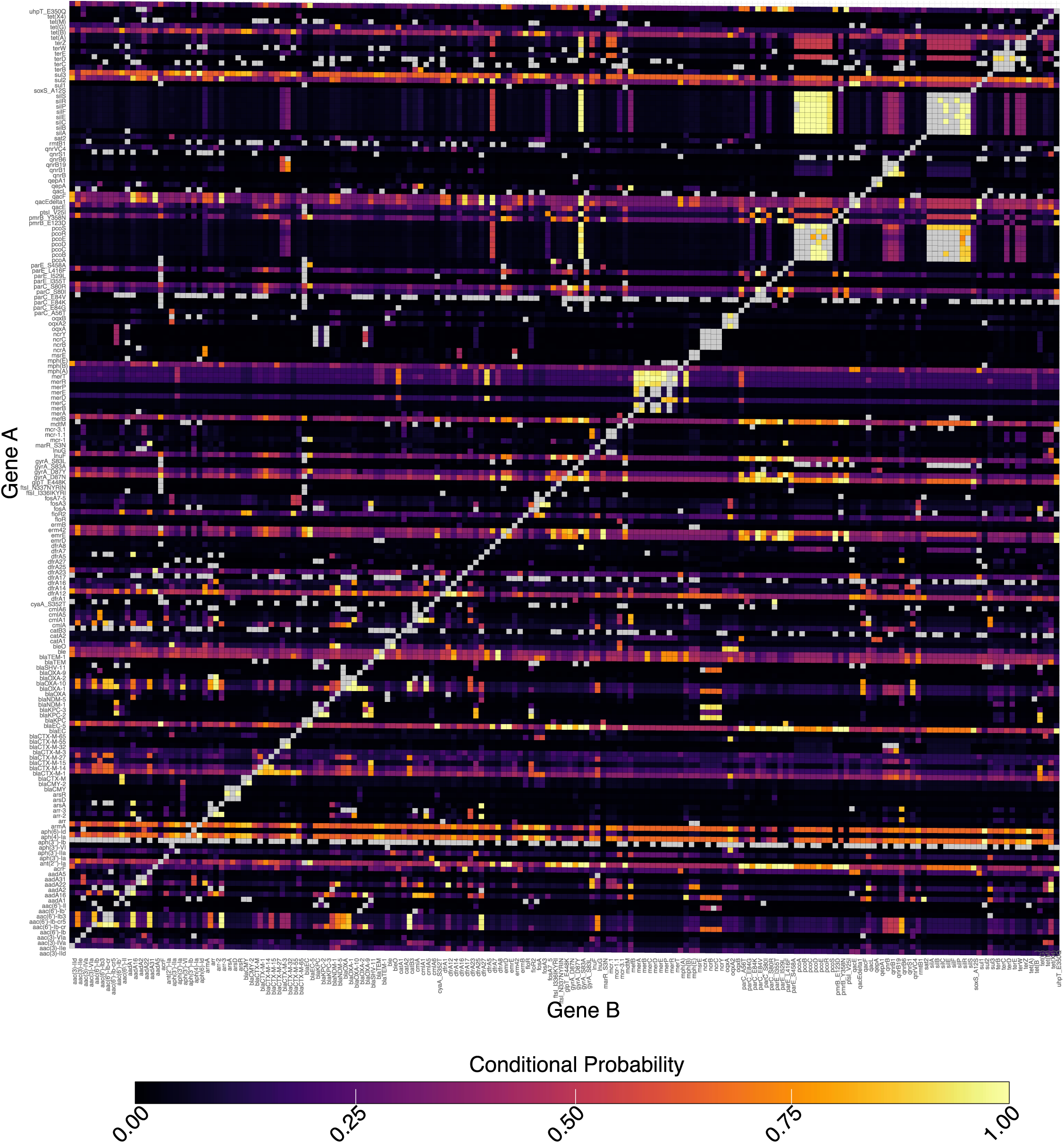
Heatmap illustrating conditional probabilities of resistance gene co-occurrence in *E. coli*. From the Bayesian network, 179 genes were queried in pairwise to calculate conditional probability of co-occurrence. Grey squares represent gene pairs excluded from the analysis due to low confidence, generation of loops in the Bayesian network, or to avoid self-query (Gene A = Gene B).

**Figure 4.**
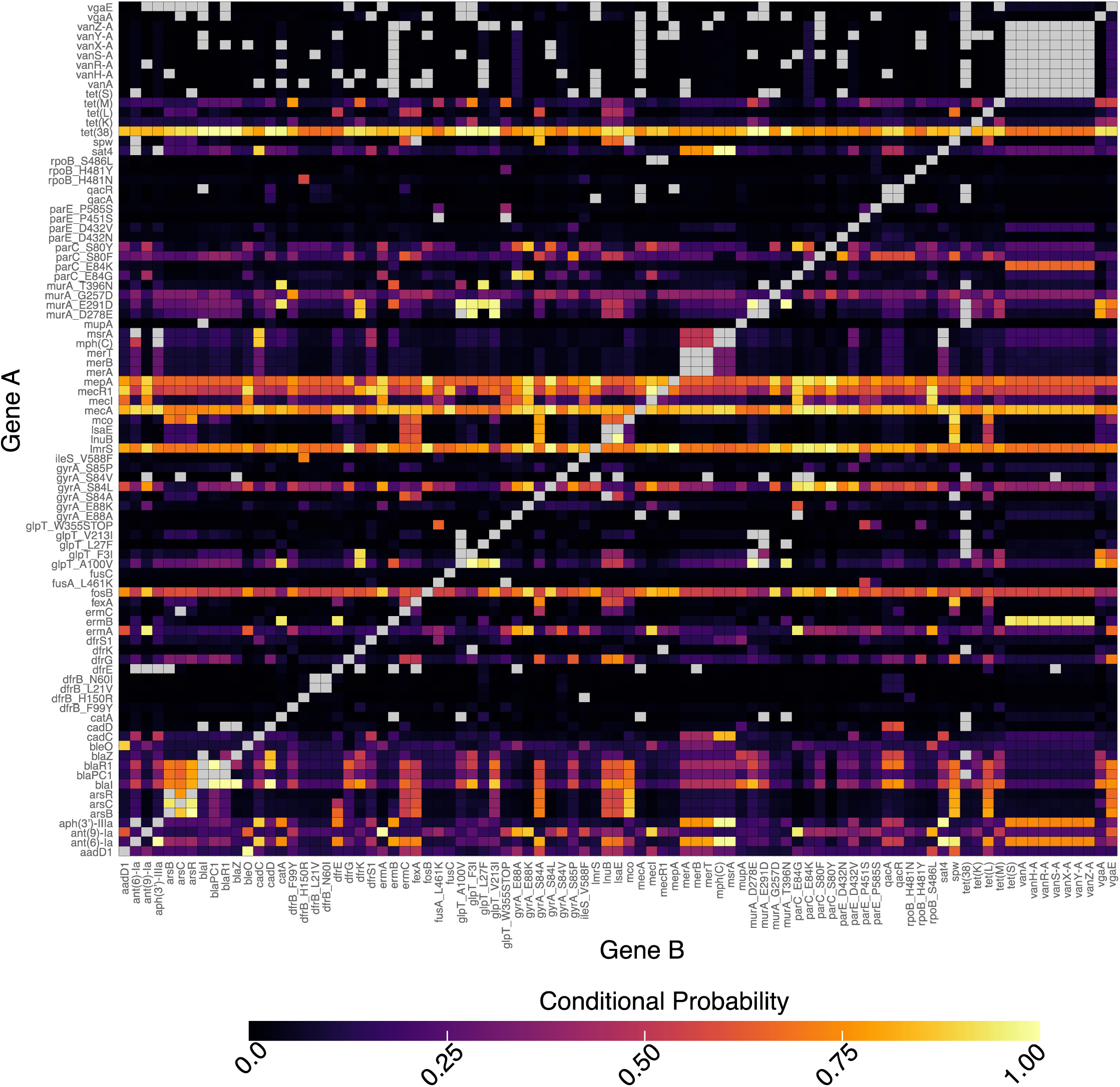
Heatmap illustrating conditional probabilities of resistance gene co-occurrence in *S. aureus*. From the Bayesian network, 91 genes were queried in pairwise to calculate conditional probability of co-occurrence. Grey squares represent gene pairs excluded from the analysis due to low confidence, generation of loops in the Bayesian network, or to avoid self-query (Gene A = Gene B).

While investigating co-occurrence patterns of resistance genes, we initially focused on the widely used trimethoprim-sulfamethoxazole, commonly employed in treating Gram-negative infections such as urinary tract infections (47,48), expecting to see strong probabilistic relationships between trimethoprim and sulfonamide resistance genes. Though not as expansive as initially predicted, co- occurrence of trimethoprim resistance-conferring dihydrofolate reductase *dfrA* genes and sulfonamide resistance *sul* genes were detected in all analyzed Gram-negative bacteria except *A. baumannii* (Table S1B and S3). Interestingly, the strongest relationship observed involving a trimethoprim resistance gene in *A. baumannii* was *dfrA1* with the streptothricin resistance-conferring N-acetyltransferase gene, *sat2* (cond. prob. 0.97, rel. risk 94.6).

Building on this, our investigation extended to patterns of resistance gene co-occurrence involving genes associated with last-line antibiotics for Gram-negative bacterial infections. A concerning observation emerged as multiple instances of ARG co-occurrence with *mcr*-family colistin resistance genes were identified in *E. coli* (Table S4). Although these relationships generally exhibited low conditional probabilities, each relationship displayed strong relative risk scores. Genes co-occurring with *mcr-1* include *tet(X4)* (cond. prob, 0.37, rel. risk 27.8, BDPS 3.2), which confers resistance to tigecycline, tetracycline resistance gene *tet(M)* (cond. prob. 0.37, rel. risk 29.2, BDPS 2.2), and streptomycin resistance gene *aadA22* (cond. prob. 0.39, rel. risk 41.6, BDPS 1.0). Of particular interest was the strong positive probabilistic relationships was observed between *mcr-1.1* and the lincosamide resistance gene *lnu(F)* (cond. prob. 0.78, rel. risk 112.0). Intriguingly, despite lincosamide’s infrequent clinical use against aerobic Gram-negative bacteria (49), this finding may indicate the acquisition of resistance genes in response to environmental pressures(50).

### Chloramphenicol Resistance Genes Exhibit Strong Co-Occurrence with Clinically Significant Resistance Genes

Several bacterial pathogens displayed positive probabilistic co-occurrence relationships with chloramphenicol resistance genes, including those associated with aminoglycoside, rifamycin, tetracycline, cephalosporin, macrolide, colistin, fosfomycin, and trimethoprim (Table 2). The most common pattern of co-occurrence involved *catB* family chloramphenicol *O*-acetyltransferases and *bla_OXA_*family cephalosporin resistance genes. *E. coli*, *K. pneumoniae*, *S. enterica*, and *P. aeruginosa* all displayed strong probabilistic relationships between these genes (Table 2) (cond. prob. 0.83-0.98, rel. risk 5.5-1187.7, BDPS 0.88-1.03), although the high relative risk observed between *catB3* and *bla_OXA-1_* in *S. enterica* could be influenced by the low occurrence of these genes (*catB3* n = 19, *bla_OXA-1_* n = 22) (Table S2). In *A. baumannii*, the strongest examples of chloramphenicol resistance gene co-occurrence included chloramphenicol resistance *O*-acetyltransferase, *catA1* (n = 26), with tetracycline resistance gene *tet(A)* (n = 18) (cond. prob. 0.93, rel. risk 96.0, BDPS 1.33) and *catB8* with aminoglycoside resistance gene *aac(6’)-Ib’* (cond. prob. 0.99, rel. risk 358.1, BDPS 1.03). Interestingly, in *A. baumannii*, *catA1* exhibited a high probability of co-occurrence with the mercury efflux gene *merE* (n = 25) (51) in (cond. prob. 0.85, rel. risk 137.4, BDPS 1.12). In *E. coli*, co-occurrence of the chloramphenicol efflux gene *cmlA1* with the colistin resistance gene *mcr-3.1* gave a strong relative risk score (cond. prob. 0.44, rel. risk 27.2, BDPS 6.4); however, the strong relative risk ratio could be due to the low sample size of *mcr-3.1* within the *E. coli* dataset (*cmlA1* n = 27, *mcr-3.1* n = 7) (Table S2).

**Table 2.** Co-occurrence of chloramphenicol resistance genes in Gram-negative bacteria. Gene A column includes only genes associated with chloramphenicol resistance. Gene B are co-occurring genes and the antibiotics they confer resistance to are listed in the Gene B Class column. ND, not determined.

| Organism | Gene A | Gene B | Gene B Antibiotic Class | Conditional Probability | SD | Relative Risk | SD | BDPS |
| --- | --- | --- | --- | --- | --- | --- | --- | --- |
| *E. coli* | *catB3* | *aac(6')-Ib-cr* | Aminoglycoside/Quinolone | 0.87 | 0.03 | 2670.67 | 1549.47 | 0.88 |
| *E. coli* | *catB3* | *bla<sub>OXA-1</sub>* | Cephalosporin | 0.83 | 0.03 | 275.72 | 292.70 | 0.88 |
| *E. coli* | *catB3* | *aac(3)-Ile* | Aminoglycoside | 0.71 | 0.05 | 16.63 | 2.31 | 1.59 |
| *E. coli* | *cmlA* | *arr_2* | Rifamycin | 0.96 | 0.03 | 43.79 | 8.25 | ND |
| *E. coli* | *cmlA* | *bla<sub>OXA-10</sub>* | Cephalosporin | 0.90 | 0.08 | 43.82 | 9.76 | ND |
| *E. coli* | *cmlA1* | *mefB* | Macrolide | 0.83 | 0.07 | 63.29 | 15.03 | 3.61 |
| *E. coli* | *cmlA1* | *aac(6')-Ib3* | Aminoglycoside | 0.78 | 0.14 | 60.90 | 18.31 | 2.89 |
| *E. coli* | *cmlA1* | *sul3* | Sulfonamide | 0.75 | 0.08 | 260.45 | 162.76 | 0.94 |
| *E. coli* | *cmlA1* | *mcr-3.1* | Polymyxin | 0.44 | 0.21 | 27.23 | 14.85 | 6.37 |
| *A. baumannii* | *catA1* | *tetA* | Tetracycline | 0.93 | 0.06 | 95.98 | 50.50 | 1.33 |
| *A. baumannii* | *catA1* | *merE* | Mercury transporter | 0.85 | 0.07 | 137.36 | 238.58 | 1.12 |
| *A. baumannii* | *catB8* | *aac(6')-Ib'* | Aminoglycoside | 0.99 | 0.00 | 358.13 | 663.16 | 1.03 |
| *A. baumannii* | *cmlA* | *arr-2* | Rifamycin | 0.93 | 0.03 | 630.26 | 1266.67 | 1.26 |
| *K. pneumoniae* | *catB* | *rmtF1* | Aminoglycoside | 0.97 | 0.03 | 20.83 | 2.46 | ND |
| *K. pneumoniae* | *catB* | *aac(6')-Ib-cr5* | Aminoglycoside/Quinolone | 0.96 | 0.02 | 155.38 | 56.92 | 1.10 |
| *K. pneumoniae* | *catB* | *bla<sub>OXA-1</sub>* | Cephalosporin | 0.95 | 0.02 | 268.00 | 107.65 | 1.03 |
| *K. pneumoniae* | *catB3* | *arr-3* | Rifamycin | 0.84 | 0.12 | 20.25 | 3.68 | ND |
| *K. pneumoniae* | *catB3* | *tetA* | Tetracycline | 0.70 | 0.06 | 32.54 | 7.67 | 1.34 |
| *S. enterica* | *catA1* | *dfrA7* | Trimethoprim | 0.99 | 0.01 | 159.67 | 41.18 | ND |
| *S. enterica* | *catB3* | *bla<sub>OXA_1</sub>* | Cephalosporin | 0.85 | 0.07 | 1187.74 | 950.50 | 0.92 |
| *S. enterica* | *catB3* | *aph(4)-Ia* | Aminoglycoside | 0.75 | 0.09 | 312.47 | 255.41 | 1.00 |
| *S. enterica* | *catB3* | *aac(3)-IVa* | Aminoglycoside | 0.73 | 0.08 | 270.50 | 127.51 | 0.99 |
| *S. enterica* | *catB3* | *arr* | Rifamycin | 0.71 | 0.09 | 695.67 | 413.00 | 0.79 |
| *P. aeruginosa* | *catB* | *aph(3')-IIb* | Aminoglycoside | 0.98 | 0.00 | 4.16 | 0.96 | 0.99 |
| *P. aeruginosa* | *catB* | *bla<sub>OXA</sub>* | Beta-lactam | 0.98 | 0.00 | 5.50 | 1.46 | 0.99 |
| *P. aeruginosa* | *catB* | *fosA* | Fosfomycin | 0.98 | 0.00 | 5.85 | 1.60 | 0.98 |

### Resistance Gene Co-Occurrence in Gram-Positive Bacterial Pathogens

In addition to Gram-negative pathogens, four Gram-positive bacterial pathogens − *E. faecalis, E. faecium, S. aureus,* and *S. pneumoniae* − were analyzed (Table 1), though it is worth nothing that *S. pneumoniae* represents our smallest Bayesian network (10 nodes). To assess the accuracy of our system, the Bayesian networks were initially analyzed for the presence of previously observed co-resistance patterns, including the co-occurrence of phenicol and lincosamide resistance genes (38,52). In *S. aureus*, the phenicol resistance gene *fexA* exhibited an overall positive probability of co-occurrence with the lincosamide resistance gene lnuB (cond. prob. 0.59, rel. risk 138.9, BDPS 0.94) (Figure 4). However, these were among the lower-occurring genes identified in *S. aureus* (*fexA* n = 23; *lnuB* n = 24) (Table S2). Among Gram-positive bacteria, *fexA* exhibited positive probabilistic relationships with genes associated with oxazolidinone (*optrA, E. faecalis*; cond. prob. 0.63, rel. risk 55.1, BDPS 0.89), aminoglycoside (*ant*(*9*)*-Ia E. faecalis*; cond. prob. 0.62, rel. risk 9.8, BDPS 1.2), and quinolone (*gyrA_S84A_, S. aureus*; cond. prob. 0.74, rel. risk 121.6, BDPS 1.5) resistance. We observed frequent and strong co-occurrence between the macrolide resistance rRNA adenine methyltransferase genes *erm(A)* and *erm(B)* with resistance genes associated with multiple classes of antibiotics. These co-occurring genes included streptomycin resistance gene *ant(6)-Ia* (*E. faecalis*; cond. prob. 0.95, rel. risk 20.3, BDPS 1.4), lincosamide resistance gene *InuB* (*E. faecalis*; cond. prob. 0.93, rel. risk 9.6, BDPS 2.5), tetracycline resistance genes *tet(L)* and *tet(S)* (*E. faecalis*; cond. prob. 0.69, rel. risk 6.1, BDPS 2.5; *S. aureus*; cond. prob. 0.95, rel. risk 63.6), and trimethoprim resistance gene *dfrE* (*S. aureus*; cond. prob. 0.94, rel. risk 59.1). Strong positive probabilistic relationships were also observed between *erm(B)* and multiple vancomycin resistance genes in *S. aureus*, including *vanA, vanH-A, vanZ-A, vanY- A, vanS-A, vanR-A,* and *vanX-A* (cond. prob. 0.91-0.94, rel. risk 59.6-62.7) (Table S5). Interestingly, *erm(B)* (n = 34) displayed a strong relationship with chloramphenicol resistance gene *catA* (n = 9) in *S. aureus* (cond. prob. 0.94, rel. risk 66.3).

As vancomycin resistance represents a major clinical challenge, we wanted to identify genes that commonly co-occur with genes associated with its resistance. Multiple vancomycin resistance genes identified in *S. aureus*, *E. faecium*, and *E. faecalis* displayed moderate (rel. risk > 2) to strong (rel. risk > 10) positive relative risk scores, indicating positive probabilistic relationships with genes conferring resistance to the macrolides, quinolones, lincosamide, aminoglycosides, and chloramphenicol (Table S6). In *S. aureus*, co-occurring genes included *vgaE, gyrA_E88A_*and *parC_E84K_, aph(3’)-IIIa, dfrE, catA,* and *glpT_L27F_*, representing lincosamide, quinolone, aminoglycoside, trimethoprim, chloramphenicol, and fosfomycin resistance, respectively (Figure 4, Table S6). In *E. faecium*, genes that co-occur with vancomycin resistance included *mef(H), gyrA_S83I_, dfrG, sat4, ant(6)-Ia, aph(3’)-IIIa,* and *23S_G2576T_,* representing macrolide, quinolone, streptothricin, aminoglycoside, and oxazolidinone resistance, respectively. In *E. faecalis*, *gyrA_S83I_*, *fexB*, and *tet(O/W/32/O)*, representing phenicol and tetracycline resistance, were found to co-occur with multiple vancomycin resistance genes. From our analysis, each organism displayed distinct, though overlapping, patterns of resistance gene co-occurrence with vancomycin resistance genes, suggesting genetic linkage or co-selection of specific resistance genes. However, in all three Gram-positive strains, the strongest probabilistic relationships were between the vancomycin resistance genes themselves, which is unsurprising, given that genes conferring resistance to vancomycin are often clustered together (39).

### Antibiotic and Heavy Metal Resistance Genes Exhibit Strong Patterns of Co-Occurrence

As heavy metal resistance genes (HMGs) may facilitate co-selection of ARGs under environmental pressures (27), we were interested in investigating whether antibiotic and heavy metal resistance genes displayed probabilistic patterns of co-occurrence within our genomic datasets. With the Gram-negative pathogens, only *E. coli, P. aeruginosa, S. enterica, A. baumannii,* and *K. pneumoniae* encoded known heavy metal resistance genes. Importantly, moderate-to-strong relationships between every type of HMG and ARGs representing nearly every antibiotic class, including the quinolones, trimethoprim, β- lactamases, chloramphenicol, rifamycin, aminoglycosides, colistin, and tetracyclines were observed. Creation of an association network allowed for visualization of HMG-ARG co-occurrence patterns across our Gram-negative dataset (Figure 5A). The strongest positive probabilistic relationship was observed between the mercuric reductase gene, *merA* (n = 33), and streptomycin resistance gene, *aadA7* (n = 23), in *S. enterica* (cond. prob. 0.94, rel. risk 274.3). In *A. baumannii*, the mercury transporter gene *merE* exhibited strong relationships with both *catA1* (cond. prob. 0.76, rel. risk 254.2, BDPS 0.89) and the tetracycline resistance gene *tet(A)* (cond. Prob. 0.98, rel. risk 244.0, BDPS 1.2). In *E. coli*, the mercuric resistance genes *merT* and *merP* displayed strong probabilistic relationships with the trimethoprim resistance gene, *dfrA5* (cond. prob. 0.91/0.90, rel. risk 6.3/6.4, BDPS 6.7/6.5, respectively) (Figure 5A). From this analysis, patterns of co-occurrence involving multiple classes of antibiotic and heavy metal resistance genes unique to each organism were detected, exposing a complex interplay between these gene categories (Figure 5B).

**Figure 5.**
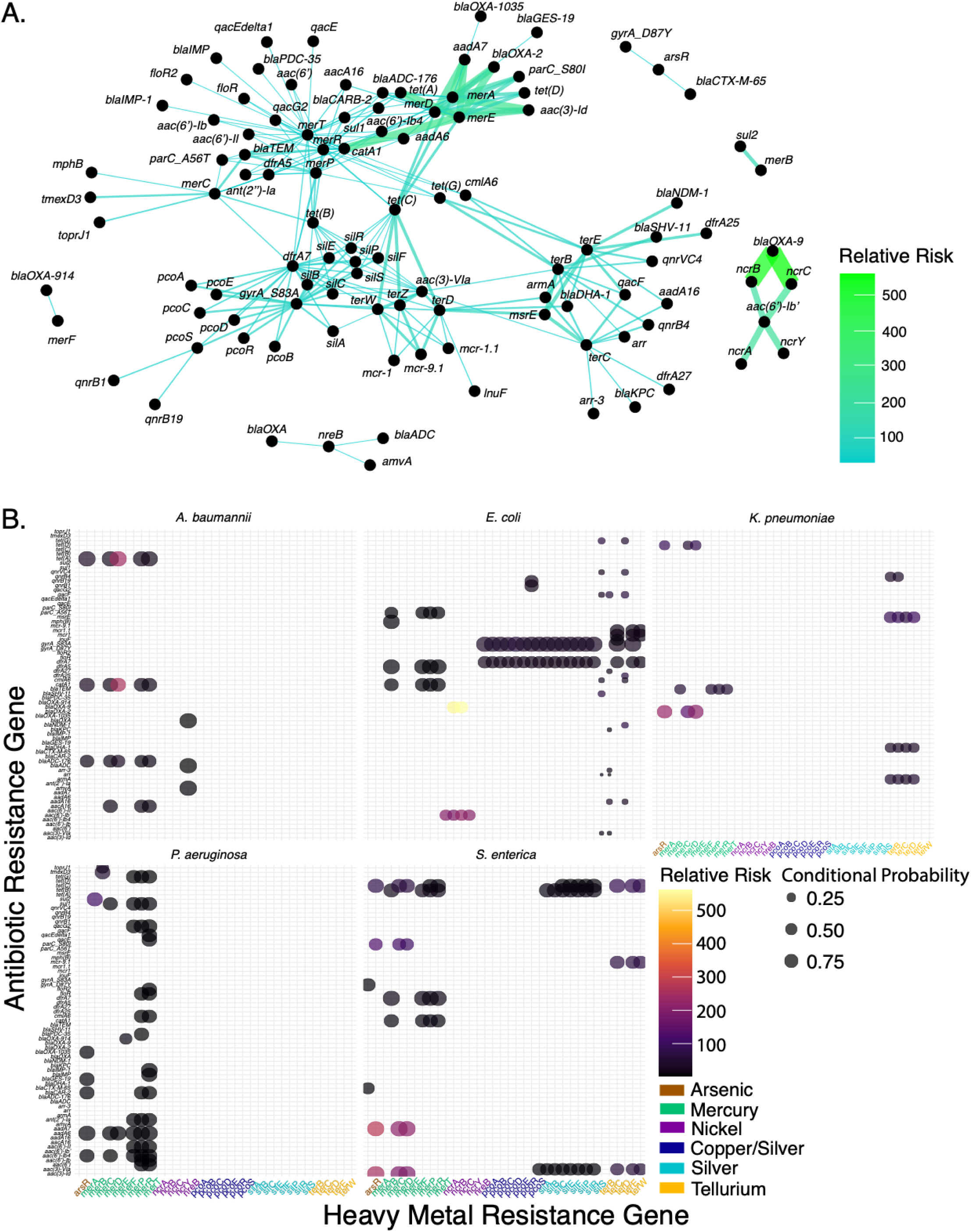
Antibiotic and heavy metal resistance gene co-occurrence in *Escherichia coli, Salmonella enterica, Pseudomonas aeruginosa, Klebsiella pneumoniae,* and *Acinetobacter. baumannii.* **A.** Association network of patterns of antibiotic and heavy metal resistance gene co-occurrence. Edge width and color is scaled by magnitude of relative risk **B.** Bubble plot illustrating antibiotic and heavy metal gene co-occurrence by organism. Bubble size is scaled by conditional probability and color is scaled by relative risk.

Among the Gram-positive pathogens analyzed, only *S. aureus* demonstrated a notable co- occurrence of HMGs and ARGS. This included genes associated with resistance to mercury, cadmium, arsenic, and copper, as well as ARGs spanning multiple classes (Figure 6A). The strongest of these relationships involved the cadmium resistance gene, *cadC*, and macrolide resistance genes *msrA* and *mph(C)* (cond. prob. 0.84/0.84, rel. risk 142.2/138.8, BDPS 0.96/0.96, respectively) (Figure 6B). Additional co-occurring ARGs included those associated with β-lactamases, streptothricin, aminoglycosides, ammonium efflux, quinolones, tetracyclines, lincosamide/streptogramin, phenicols, and fosfomycin, respectively (Figure 6). In *E. faecalis,* two gene pairs were detected, involving cadmium and mercury, as well as lincosamide and β-lactamase resistance genes. Despite having low conditional probabilities, these pairs exhibited strong relative risk values (Figure 6B).

**Figure 6.**
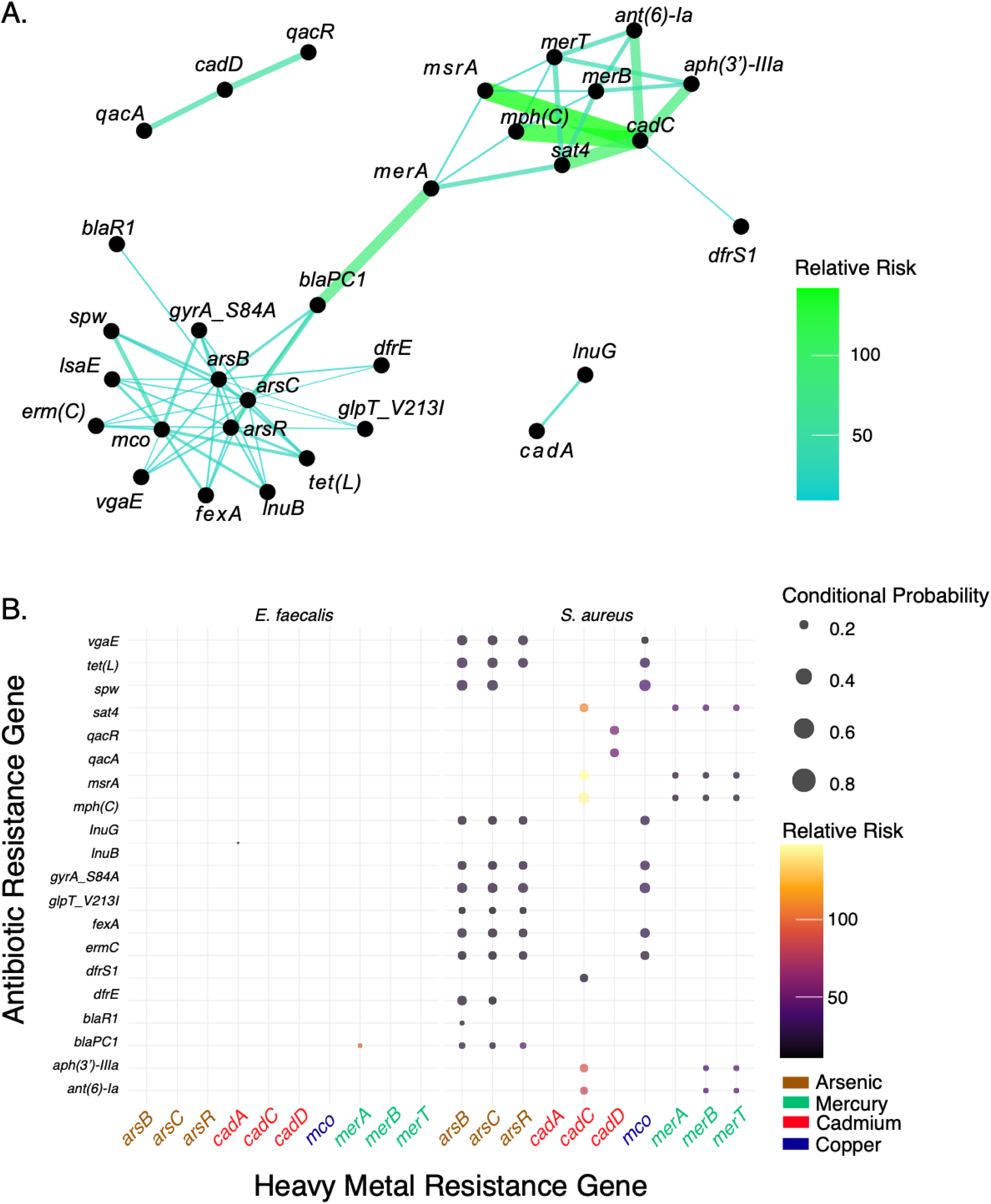
Antibiotic and heavy metal resistance gene co-occurrence in *Enterococcus faecalis* and *Staphylococcus aureus.* **A.** Association network of patterns of antibiotic and heavy metal resistance gene co-occurrence. Edge width and color is scaled by relative risk **B.** Bubble plot illustrating antibiotic and heavy metal gene co-occurrence by organism. Bubble size is scaled by conditional probability and color is scaled by relative risk.

## Discussion

Rising multidrug resistance rates represent a global health threat that is unlikely to be solved anytime soon (1,5,6,8,13). However, the effective utilization of probabilistic statistical models can offer some predictive value to trends and patterns of resistance gene co-occurrence. The development and application of the ReGAIN bioinformatic pipeline, which leverages a wealth of publicly available genomic data, serves to offer a reproducible method of measuring resistance gene co-occurrence. Additionally, ReGAIN may facilitate the identification of previously unrecognized gene cohorts that work synergistically or provide avenues for resistance gene co-selection under clinical or environmental pressures (29). By identifying these relationships, genes can be prioritized for further characterization. In this study, several interesting probabilistic relationships were identified from Gram-negative and Gram-positive bacterial pathogens, including several unexpected relationships between clinically important ARGs and various chloramphenicol resistance genes (Table 2). Although chloramphenicol is infrequently used in clinical human health settings in the United States (53), the NCBI database contains genomes deposited from all over the world. Thus, it is plausible that our datasets contain genomes isolated from patients in other countries, which could explain the strong relationships observed. Alternatively, we observed similar relationships in a genomic population of *E. coli* isolated from hospitalized patients in Salt Lake City, Utah (unpublished data). This suggests that the co-occurrence of chloramphenicol and other resistance genes could be evidence of the mobilization of MDR strains through the food chain, as bacteria harboring phenicol resistance genes have been identified in food animals (46,54).

Besides mapping common patterns of resistance gene co-occurrence, we were additionally interested in investigating ARG co-occurrence involving problematic resistance genes, such as those conferring resistance to vancomycin, methicillin, or last-line antibiotics like colistin. Though several *mcr-* family colistin resistance genes have been observed occurring together (55), to the best of our knowledge, this study is the first to explore co-occurrence of *mcr-*family genes with other antibiotic resistance genes using a large-scale genomic analysis. Indeed, several potentially problematic relationships involving colistin resistance genes were identified, including co-occurrence with tetracycline, sulfonamide, and aminoglycoside resistance genes.

Multidrug resistance is an ever-evolving threat, and it is important that trends of gene co-occurrence are identified and cataloged. Among resistance genes, relationships between antibiotic and heavy metal resistance genes are still not fully understood, though there is evidence of synergy or co-regulation between these gene categories (28,29,56). Surprisingly, this study identified multiple examples of ARG-HMG co-occurrence in both Gram-negative and Gram-positive bacterial pathogens (Figures 5 and 6). Moreover, several ARG-HMG gene pairs displayed BDPS scores indicating that the presence of the HMG was influenced by the presence of the ARG (Table S7). Examples include the mercury resistance gene *merT* and the carbapenem resistance β-lactamase gene *bla_IMP-1_* in *P. aeruginosa* (BDPS 14.3); the silver resistance gene *silC* and gentamicin resistance gene *aac*(*3*)*-Via* in *S. enterica* (BDPS 32.6); and in *S. aureus*, the copper resistance gene *mco* and lincosamide resistance gene *vgaE* (BDPS 13.9) as well as the arsenic resistance gene *arsC* and trimethoprim resistance gene *dfrE* (BDPS 13.2). In each case, the probability of observing the HMG was 13 to 32-fold higher given the presence of the ARG. Conversely, multiple examples were identified where the probability of observing the ARG was higher given the presence of the HMG, indicating that the presence of the ARG may be influenced by presence of the HMG (Table S7). Taken together, these results could indicate co-selection of specific heavy metal and antibiotic resistance genes. By using relative risk (rel. risk < 1 indicates a higher probability of observing Gene A in the absence of Gene B), we can also identify antagonistic relationships, revealing patterns of genes that either cannot or do not co-occur.

Though Bayesian network structure learning represents a powerful statistical approach to identifying and mapping co-occurrence of resistance genes, it is important to understand the limitations of these models. Sample size and isolation source can skew results, especially if sample size is very small or isolation source is biased towards one environment. To ensure an accurate overview of organism-specific patterns of resistance, it is important that large sample sizes representing myriad isolation sources, including clinical, environmental, and agricultural are used. Furthermore, to track movement of resistance genes, the isolation source, origin, and year of collection should be documented when genomic data are deposited. Using well-curated genomic databases will help facilitate tracking the spread of resistance genes in a region and time-specific manner.

Overall, the utilization of the ReGAIN platform offers a broad look at common patterns of antibiotic and heavy metal resistance gene co-occurrence in clinically relevant and high threat bacterial pathogens. By using large genomic populations pulled from publicly available databases, this study works to mitigate potential bias introduced by oversampling from either clinical or environmental sources, which allows for general inferences to be made concerning commonly co-occurring resistance genes. Furthermore, making the ReGAIN bioinformatic pipeline available as both a web interface and command line software provides researchers with a reproducible probabilistic statistical method of measuring resistance gene co-occurrence.

## Methods

### Data Acquisition

All genomic data was downloaded from the National Center for Biotechnology Information (NCBI) using either the NCBI web server or the NCBI Datasets command line software (57). To ensure only full genomes or large plasmid genomic data was included in each analysis, FASTA files containing < 3500 nucleotides were filtered and excluded using a custom Python script. Antibiotic resistance genes were identified using AMRfinderPlus v.3.10.40 (32) using the appropriate organism flag. The ReGAIN data acquisition workflow was built using Python v.3.10.12.

### Bayesian Analysis, and Results Visualization

The Bayesian network structure learning module was built using R v.4.3.1 and the following packages: bnlearn v.4.8.1 (40), gRain v.1.3.13 (41), gRbase v.1.8.9 (58), visNetwork v.2.1.2, and igraph v.1.5.0 (59). Results were visualized using ggplot2 v.3.4.2 (60). Bayesian Networks were built using the bnlearn and gRain packages using 500 bootstraps; data was additionally resampled and fitted to networks 100 times to calculate confidence. Laplace smoothing was implemented that scaled with the number of variables to avoid divide by zero errors while calculating relative risk. Given *N* variables, conditional probabilities were adjusted by [(*N* + 0.5) / (*N* + 1)], which maintains the appropriate scale while avoiding *inf* values. For each genomic dataset, only genes occurring ≥ 5 times were included. PCoA was created using the Jaccard method of distance and the following R packages: ellipse v.0.3.2 and vegan v.2.6-4 (61). To reduce noise in the network, genes occurring less than five times across within each genomic population were not included in the analysis. Antibiotic and heavy metal resistance gene association networks were created using ggraph v.2.1.0 (62). Not determined (ND) values indicate that a gene pair was excluded from the analysis due to low confidence (< 0.5) or because the pair would introduce cycles in the Bayesian network.

## Author Contributions

**Elijah R. Bring Horvath:** conceptualization (equal); bioinformatics (lead); programming (lead); data analysis (lead); writing – original draft (lead); writing – review and editing (equal). **Mathew G. Stein:** programming (equal); writing – review and editing (equal). **Matthew Mulvey:** conceptualization (equal); data analysis (equal); writing − review and editing (equal). **Edgar Javier Hernandez:** conceptualization (equal); bioinformatics (equal); data analysis (equal); writing – original draft (equal); writing – review and editing (equal). **Jaclyn M. Winter**: conceptualization (equal); project administration (lead); data analysis (equal); writing − original draft (equal); writing − review and editing (equal).

## Data Availability Statement

The ReGAIN command line software is available at https://github.com/ERBringHorvath/regain_cl.

## Other

ReGAIN is under provisional patent US 63/526,656.

**Equation 1.** Conditional probability. The probability of observing ‘Gene A’ given the presence of ‘Gene B’.

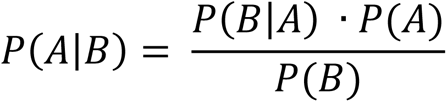

**Equation 2.** Relative risk. The ratio of the conditional probability of observing ‘Gene A’ given the presence of ‘Gene B’ to the conditional probability of observing ‘Gene A’ in the absence of ‘Gene B’. This calculation adjusts for single occurrence of Gene A in the dataset, offering a more in-depth understanding of the probability and rate of co-occurrence.

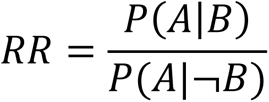

**Equation 3.** Bidirectional probability score. The ratio of the conditional probability of observing ‘Gene A’ given the presence of ‘Gene B’ to the conditional probability of observing ‘Gene B’ given the presence of ‘Gene A’.

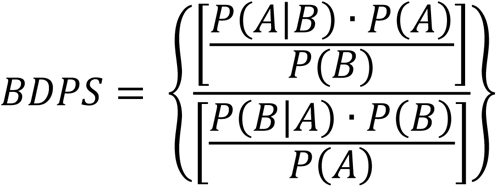

**Equation 4.** Fold change. The ratio of the relative risk of ‘Gene A’ to ‘Gene B’ to the relative risk of ‘Gene B’ to ‘Gene A’.

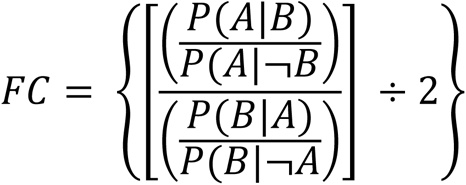

## References

1. Prestinaci F, Pezzotti P, Pantosti A. (2015) Antimicrobial resistance: a global multifaceted phenomenon. Pathog Glob Health, 109, 309–318.

2. Klein EY, Tseng KK, Pant S, Laxminarayan R. (2019) Tracking global trends in the effectiveness of antibiotic therapy using the Drug Resistance Index. BMJ Glob Health, 4, e001315.

3. 3. Centers for Disease Control and Prevention, U.D.o.H.a.H.S. (2021). Centers for Disease Control and Prevention, National Center for Emerging and Zoonotic Infectious Diseases (NCEZID), Division of Healthcare Quality Promotion (DHQP), Vol. 2023.

4. Centers for Disease Control and Prevention US Department of Health and Human Services. (2019) Antibiotic Resistance Threats in the United States. U.S. Department of Health and Human Services, CDC.

5. Sharma S, Barman P, Joshi S, Preet S, Saini A. (2022) Multidrug resistance crisis during COVID-19 pandemic: Role of anti-microbial peptides as next-generation therapeutics. Colloids Surf B Biointerfaces, 211, 112303.

6. Munita JM, Arias CA. (2016) Mechanisms of Antibiotic Resistance. Microbiol Spectr, 4.

7. Forde BM, Roberts LW, Phan MD, Peters KM, Fleming BA, Russell CW, Lenherr SM, Myers JB, Barker AP, Fisher MA, Chong TM, Yin WF, Chan KG, Schembri MA, Mulvey MA, Beatson SA. (2019) Population dynamics of an Escherichia coli ST131 lineage during recurrent urinary tract infection. Nat Commun, 10, 3643.

8. Diekema DJ, Hsueh PR, Mendes RE, Pfaller MA, Rolston KV, Sader HS, Jones RN. (2019) The Microbiology of Bloodstream Infection: 20-Year Trends from the SENTRY Antimicrobial Surveillance Program. Antimicrob Agents Chemother, 63, e00355–00319.

9. Ikhimiukor OO, Oaikhena AO, Afolayan AO, Fadeyi A, Kehinde A, Ogunleye VO, Aboderin AO, Oduyebo OO, Elikwu CJ, Odih EE, Komolafe I, Argimon S, Egwuenu A, Adebiyi I, Sadare OA, Okwor T, Kekre M, Underwood A, Ihekweazu C, Aanensen DM, Okeke IN. (2022) Genomic characterization of invasive typhoidal and non-typhoidal Salmonella in southwestern Nigeria. PLoS Negl Trop Dis, 16, e0010716.

10. Hutchings MI, Truman AW, Wilkinson B. (2019) Antibiotics: past, present and future. Curr Opin Microbiol, 51, 72–80.

11. Nandi A, Peceta S, Bloom DE. (2023) Global antibiotic use during the COVID-19 pandemic: analysis of pharmaceutical sales data from 71 countries, 2020-2022. EClinicalMedicine, 57, 101848.

12. Machowska A, Lundborg CS. (2018) Drivers of Irrational Use of Antibiotics in Europe. Int J Environ Res Public Health, 16, 27.

13. Manyi-Loh C, Mamphweli S, Meyer E, Okoh A. (2018) Antibiotic Use in Agriculture and Its Consequential Resistance in Environmental Sources: Potential Public Health Implications. Molecules, 23, 795.

14. Bischoff, K.M., White, D.G., Hume, M.E., Poole, T.L. and Nisbet, D.J. (2005) The chloramphenicol resistance gene cmlA is disseminated on transferable plasmids that confer multiple-drug resistance in swine Escherichia coli. FEMS Microbiol Lett, 243, 285–291.

15. de Vries, L.E., Valles, Y., Agerso, Y., Vaishampayan, P.A., Garcia-Montaner, A., Kuehl, J.V., Christensen, H., Barlow, M. and Francino, M.P. (2011) The gut as reservoir of antibiotic resistance: microbial diversity of tetracycline resistance in mother and infant. PLoS One, 6, e21644.

16. Hussein NH, Al-Kadmy IMS, Taha BM, Hussein JD,. (2021) Mobilized colistin resistance (mcr) genes from 1 to 10: a comprehensive review. Mol Biol Rep, 48, 2897–2907.

17. 17. Lambrecht E, Van Coillie E, Van Meervenne E, Boon N, Heyndrickx M, Van de Wiele T. (2019) Commensal E. coli rapidly transfer antibiotic resistance genes to human intestinal microbiota in the Mucosal Simulator of the Human Intestinal Microbial Ecosystem (M-SHIME). Int J Food Microbiol, 311, 108357.

18. Smillie, C.S., Smith, M.B., Friedman, J., Cordero, O.X., David, L.A. and Alm, E.J. (2011) Ecology drives a global network of gene exchange connecting the human microbiome. Nature, 480, 241–244.

19. Stecher, B., Denzler, R., Maier, L., Bernet, F., Sanders, M.J., Pickard, D.J., Barthel, M., Westendorf, A.M., Krogfelt, K.A., Walker, A.W. et al. (2012) Gut inflammation can boost horizontal gene transfer between pathogenic and commensal Enterobacteriaceae. Proc Natl Acad Sci U S A, 109, 1269–1274.

20. von Wintersdorff, C.J., Penders, J., van Niekerk, J.M., Mills, N.D., Majumder, S., van Alphen, L.B., Savelkoul, P.H. and Wolffs, P.F. (2016) Dissemination of Antimicrobial Resistance in Microbial Ecosystems through Horizontal Gene Transfer. Front Microbiol, 7, 173.

21. Bogri A, Jensen EEB, Borchert AV, Brinch C, Otani S, Aarestrup FM. (2023) Transmission of antimicrobial resistance in the gut microbiome of gregarious cockroaches: the importance of interaction between antibiotic exposed and non-exposed populations. mSystems, Advance online publication, e0101823.

22. Poole, T.L., Callaway, T.R., Bischoff, K.M., Warnes, C.E. and Nisbet, D.J. (2006) Macrolide inactivation gene cluster mphA-mrx-mphR adjacent to a class 1 integron in Aeromonas hydrophila isolated from a diarrhoeic pig in Oklahoma. J Antimicrob Chemother, 57, 31–38.

23. Huyan, J., Tian, Z., Zhang, Y., Zhang, H., Shi, Y., Gillings, M.R. and Yang, M. (2020) Dynamics of class 1 integrons in aerobic biofilm reactors spiked with antibiotics. Environ Int, 140, 105816.

24. Navas J, Fernandez-Martinez M, Salas C, Cano ME, Martinez-Martinez L. (2016) Susceptibility to Aminoglycosides and Distribution of aph and aac(3)-XI Genes among Corynebacterium striatum Clinical Isolates. PLoS One, 11, e0167856.

25. Baker M, Zhang X, Maciel-Guerra A, Babaarslan K, Dong Y, Wang W, Hu Y, Renney D, Liu L, Li H, Hossain M, Heeb S, Tong Z, Pearcy N, Zhang M, Geng Y, Zhao L, Hao Z, Senin N, Chen J, Peng Z, Li F, Dotorini T. (2024) Convergence of resistance and evolutionary responses in Escherichia coli and Salmonella enterica co-inhabiting chicken farms in China. Nat Commun, 15, 206.

26. D’Costa VM, King CE, Kalan L, Morar M, and Sung WW, S.C., Froese D, Zazula G, Calmels F, Debruyne R, Golding GB, Poinar HN, Wright GD. (2011) Antibiotic resistance is ancient. Nature, 477, 457–461.

27. Li LG, Xia Y, Zhang T. (2017) Co-occurrence of antibiotic and metal resistance genes revealed in complete genome collection. ISME J, 11, 651–662.

28. Akinbowale OL, Peng H, Grant P, Barton MD. (2007) Antibiotic and heavy metal resistance in motile aeromonads and pseudomonads from rainbow trout (Oncorhynchus mykiss) farms in Australia. Int J Antimicrob Agents, 30, 177–182.

29. Zhang H, Ma Y, Liu P, Li X. (2016) Multidrug resistance operon emrAB contributes for chromate and ampicillin co-resistance in a Staphylococcus strain isolated from refinery polluted river bank. Springerplus, 5, 1648.

30. Liu YY, Wang Y, Walsh TR, Yi LX, Zhang R, Spencer J, Doi Y, Tian G, Dong B, Huang X, Yu LF, Gu D, Ren H, Chen X, Lv L, He D, Zhou H, Liang Z, Liu JH, Shen J. (2016) Emergence of plasmid-mediated colistin resistance mechanism MCR-1 in animals and human beings in China: a microbiological and molecular biological study. Lancet Infect Dis, 16, 161–168.

31. Ordooei JA, Shokouhi S, Sahraei Z. (2015) A review on colistin nephrotoxicity. Eur J Clin Pharmacol, 71, 801–810.

32. Feldgarden M, Brover V, Gonzalez-Escalona N, Frye JG, Haendiges J, Haft DH, Hoffmann M, Petengill JB, Prasad AB, Tillman GE, Tyson GH, Klimke W. (2021) AMRFinderPlus and the Reference Gene Catalog facilitate examination of the genomic links among antimicrobial resistance, stress response, and virulence. Sci Rep, 11, 12728.

33. Florensa AF, Kaas RS, Clausen PTLC, Aytan-Aktug D, Aarestrup FM. (2022) ResFinder - an open online resource for identification of antimicrobial resistance genes in next-generation sequencing data and prediction of phenotypes from genotypes. Microb Genom, 8, 000748.

34. Shinde, V. (2022), University of Georgia, University of Georgia.

35. Li B, Yang Y, Ma L, Ju F, Guo F, Tiedje JM, Zhang T. (2015) Metagenomic and network analysis reveal wide distribution and co-occurrence of environmental antibiotic resistance genes. ISME J, 9, 2490–2502.

36. Dhariwal A, Junges R, Chen T, Petersen FC. (2021) ResistoXplorer: a web-based tool for visual, statistical and exploratory data analysis of resistome data. NAR Genom Bioinform, 3, lqab018.

37. Szczepanowski, R., Krahn, I., Linke, B., Goesmann, A., Puhler, A. and Schluter, A. (2004) Antibiotic multiresistance plasmid pRSB101 isolated from a wastewater treatment plant is related to plasmids residing in phytopathogenic bacteria and carries eight different resistance determinants including a multidrug transport system. Microbiology (Reading), 150, 3613–3630.

38. Shen W, Zhang R, Cai J. (2023) Co-occurrence of multiple plasmid-borne linezolid resistance genes-optrA, cfr, poxtA2 and cfr(D) in an Enterococcus faecalis isolate from retail meat. J Antimicrob Chemother, 78, 1637–1643.

39. Selim S. (2022) Mechanisms of gram-positive vancomycin resistance (Review). Biomed Rep, 16, 7.

40. Scutari M. (2010) Learning Bayesian Networks with the bnlearn R Package. Journal of Statistical Software, 35, 1–20.

41. Højsgaard, S. (2012) Graphical Independence Networks with the gRain Package for R. Journal of Statistical Software, 46, 1–26.

42. Warne DJ, Baker RE, Simpson MJ. (2020) A practical guide to pseudo-marginal methods for computational inference in systems biology. J Theor Biol, 496, 110255.

43. Ko Y, Kim J, Rodriguez-Zas SL. (2019) Markov chain Monte Carlo simulation of a Bayesian mixture model for gene network inference. Genes Genomics, 41, 547–555.

44. Murdoch DJ, Chow ED. (1996) A Graphical Display of Large Correlation Matrices. The American Statistician, 50, 178–180.

45. Tang B, Chang J, Chen Y, Lin J, Xiao X, Xia X, Lin J, Yang H, Zhao G. (2022) Escherichia fergusonii, an Underrated Repository for Antibicrobial Resistance in Food Animals. Microbiology Spectrum, 10, e0161721.

46. Indugu N, Sharma L, Jackson CR, Singh P. (2020) Whole-Genome Sequence Analysis of Multidrug- Resistant *Enterobacter hormaechei* Isolated from Imported Retail Shrimp. Microbiology Resource Announcements, 9, e01103–01120.

47. Cockerill FR, Edson RS. (1991) Trimethoprim-sulfamethoxazole. Mayo Clinic proceedings, 66, 1260–1269.

48. Theibault T. (2020) Sulfamethoxazole/trimethoprim ratio as a new marker in raw wastewaters: a critical review. Science of the Total Environment, 715, 136916.

49. Matzo D, Eyal Z, Benhamou RI, Shalev-Benami M, Halfon Y, Krupkin M, Zimmerman E, Rozenberg H, Bashan A, Fridman M, Yonath A. (2017) Structural insights of lincosamides targeting the ribosome of Staphylococcus aureus. Nucleic Acids Res, 45, 10284–10292.

50. Chaturvedi P, Singh A, Chowdhary P, Pandey A, Gupta P. (2021) Occurrence of emerging sulfonamide resistance (sul1 and sul2) associated with mobile integrons-integrase (intI1 and intI2) in riverine systems. Sci Total Environ, 751, 142217.

51. Kiyono M, Sone Y, Nakamura R, Pan-Hou H, Sakabe K. (2009) The MerE protein encoded by transposon Tn*21* is a broad mercury transporter in *Escherichia coli*. FEBS Letters, 583, 1127–1131.

52. Ruekit S, Srijan A, Serichantalergs O, Margulieu KR, Mc Gann P, Mills EG, Stribling WC, Pimsawat T, Kormanee R, Nakornchai S, Sakdinava C, Sukhchat P, Wojnarski M, Demons ST, Crawford JM, Lertsethtakarn P, Swierczewski BE. (2022) Molecular characterization of multidrug-resistant ESKAPEE pathogens from clinical samples in Chonburi, Thailand (2017-2018). BMC Infect Dis, 22, 695.

53. Oong GC, Tadi P. (2023) Chloramphenicol. *StatPearls, National Library for Biotechnology Information*, Web Article.

54. Sharma L, Nagpal R, Jackson CR, Patel D, Singh P. (2021) Antibiotic-resistant bacteria and gut microbiome communities associated with wild-caught shrimp from the United States versus imported farm-raised retail shrimp. Sci Rep, 11, 3356.

55. Hernandez M, Iglesias MR, Rodriguez-Lazaro D, Gallardo A, Quijada NM, Miguela-Villoldo P, Campos MJ, Piriz S, Lopez-Orozco G, de Frutos C, Saez JL, Ugarte-Ruiz M, Dominguez L, Quesada A. (2017) Co-occurrence of colistin-resistance genes *mcr-1* and *mcr-3* among multidrug-resistant *Escherichia coli* isolated from catle, Spain, September 2015. Euro Surveill, 22, 30586.

56. Yang S, Deng W, Liu S, Yu X, Mustafa GR, Chen S, He L, Ao X, Yang Y, Zhou K, Li B, Han X, Xu X, Zou L. (2020) Presence of heavy metal resistance genes in Escherichia coli and Salmonella isolates and analysis of resistance gene structure in E. coli E308. J Glob Antimicrob Resist, 21, 420–426.

57. Sayers EW, Bolton EE, Brister JR, Canese K, Chan J, Comeau DC, Connor R, Funk K, Kelly C, Kim S, Madej T, Marchler-Bauer A, Lanczycki C, Lathrop S, Lu Z, Thibaud-Nissan F, Murphy T, Phan L, Skripchenko Y, Tse T, Wang J, Williams R, Trawick BW, Pruit K, Sherry ST. (2022) Database resources of the national center for biotechnology information. Nucleic Acids Res, 7, D20–D26.

58. Dethlefsen C, Højsgaard S. (2005) A Common Platform for Graphical Models in R: The gRbase Package. Journal of Statistical Software, 14, 1–12.

59. Csardi G, Nepusz T. (2006) The igraph software package for complex network research. *InterJournal*, Complex Systems, 1695, 1–9.

60. Wickham H. (2016) ggplot2: Elegant Graphics for Data Analysis. Springer-Verlog New York.

61. Oksanen J, Simpson G, Blanchet F, Kindt R, Legendre P, Minchin P, O’Hara R, Solymos P, Stevens M, Szoecs E, Wagner H, Barbour M, Bedward M, Bolker B, Borcard D, Carvalho G, Chirico M, De Caceres M, Durand S, Evangelista H, FitzJohn R, Friendly M, Furneaux B, Hannigan G, Hill M, Lahti L, McGlinn D, Ouellete M, Ribeiro Cunha E, Smith T, Stier A, Ter Braak C, Weedon J. (2022) vegan: Community Ecology Package.

62. Pedersen TL. (2022), https://ggraph.data-imaginist.com, https://github.com/thomasp85/ggraph.

