## Supplementary material for "Resistance Gene Association and Inference Network (ReGAIN): A Bioinformatics Pipeline for Assessing Probabilistic Co-Occurrence Between Resistance Genes in Bacterial Pathogens": SI

**Author Information**

Elijah R. Bring Horvath^1,2^, Mathew G. Stein, Matthew A. Mulvey^3,4^, Edgar J. Hernandez^5,*^, Jaclyn M. Winter^1,*^

^1^Department of Pharmacology and Toxicology, University of Utah, Salt Lake City, Utah, 84112, United States

^2^Department of Medicinal Chemistry, University of Utah, Salt Lake City, Utah, 84112, United States

^3^School of Biological Sciences, University of Utah, Salt Lake City, UT 84112, United States

^4^Henry Eyring Center for Cell & Genome Science, University of Utah, Salt Lake City, UT 84112, United States

^5^Department of Biomedical Informatics, University of Utah, Salt Lake City, Utah, 84112, United States

**Table of Contents**

**Supplementary Tables**

Table S1 A−M: Probability values for investigated bacterial pathogens	**S3**

Table S2: Occurrence of resistance genes in each genomic population	**S4**

Table S3: Co-occurrence of trimethoprim and sulfonamide resistance genes in Gram-negative bacteria	**S5**

Table S4: Co-occurrence of colistin and other resistance genes in *Escherichia coli*	**S6**

Table S5: Common patterns of resistance gene co-occurrence in *S. aureus*	**S7**

Table S6: Co-occurrence of vancomycin and other resistance genes in *Staphylococcus aureus*	**S9**

Table S7: Co-occurrence of heavy metal and antibiotic resistance genes in Gram-negative bacteria	**S12**

**Supplementary Figures**

Figure S1: Required format of externally prepared data using ReGAIN	**S13**

Figure S2: Resistance gene classes identified in each Gram-negative genomic population	**S14**

Figure S3: Resistance gene classes identified in each Gram-positive genomic population	**S15**

Figure S4: Resistance gene classes identified in *Clostridium difficile* genomic population	**S17**

**Table S1 A−M.** Conditional probability, relative risk ratio, and confidence interval values for **A.** *Escherichia coli,* **B.** *Acinetobacter baumannii,* **C.** *Enterobacter cloacae,* **D.** *Enterococcus faecalis,* **E.** *Enterococcus faecium,* **F.** *Klebsiella pneumoniae,* **G.** *Neisseria gonorrhoeae,* **H.** *Pseudomonas aeruginosa,* **I.** *Salmonella enterica,* **J.** *Salmonella enterica* subsp. *typhi,* **K.** *Staphylococcus aureus,* **L.** *Streptococcus pneumoniae,* and **M.** *Clostridium difficile*. Due to the size of this table, it is available as thirteen separate downloadable files from https://github.com/ERBringHorvath/regain_cl/tree/main/probability_tables

**Table S2**. Gene occurrence across each genomic population. Due to the size of this table, it is available as a separate download from https://github.com/ERBringHorvath/regain_cl/tree/main/probability_tables

**Table S3.** Co-occurrence of trimethoprim (Gene A) and sulfonamide resistance genes in Gram-negative bacteria. ND = not determined, SD = standard deviation, CI = confidence interval, BDPS = bidirectional probability score.

| **Organism** | **Gene A** | **Gene B** | **Gene B Resistance Type** | **Conditional Probability** | **SD** | **CI (low)** | **CI (high)** | **Relative Risk** | **SD** | **CI (low)** | **CI (high)** | **BDPS** |
| --- | --- | --- | --- | --- | --- | --- | --- | --- | --- | --- | --- | --- |
| *E. coli* | *dfrA1* | *sul1* | Sulfonamide | 0.80 | 0.02 | 0.80 | 0.81 | 3.90 | 0.27 | 3.84 | 3.95 | 1.30 |
| *E. coli* | *dfrA12* | *sul3* | Sulfonamide | 0.47 | 0.07 | 0.46 | 0.49 | 13.42 | 2.06 | 13.01 | 13.83 | 2.30 |
| *E. coli* | *dfrA17* | *sul1* | Sulfonamide | 0.70 | 0.02 | 0.70 | 0.71 | 19.89 | 3.92 | 19.11 | 20.66 | 0.79 |
| *A. baumannii* | *dfrA1* | *sat2* | Sulfonamide | 0.97 | 0.02 | 0.96 | 0.97 | 94.55 | 36.65 | 87.27 | 101.82 | ND |
| *K. pneumoniae* | *dfrA1* | *sat2* | Sulfonamide | 0.99 | 0.01 | 0.99 | 0.99 | 28.01 | 2.91 | 27.43 | 28.58 | ND |
| *K. pneumoniae* | *dfrA1* | *sul2* | Sulfonamide | 0.56 | 0.07 | 0.55 | 0.57 | 13.73 | 2.33 | 13.26 | 14.19 | 2.07 |
| *S. enterica* | *dfrA1* | *sul3* | Sulfonamide | 0.91 | 0.06 | 0.90 | 0.92 | 58.02 | 12.32 | 55.58 | 60.47 | ND |
| *S. enterica* | *dfrA12* | *sul3* | Sulfonamide | 0.43 | 0.07 | 0.41 | 0.44 | 63.96 | 17.50 | 60.49 | 67.44 | 0.94 |
| *P. aeruginosa* | *dfrA1* | *sul1* | Sulfonamide | 0.07 | 0.02 | 0.06 | 0.07 | 23.88 | 19.12 | 20.09 | 27.68 | 0.082 |
| *P. aeruginosa* | *dfrB2* | *sul1* | Sulfonamide | 0.06 | 0.02 | 0.06 | 0.06 | 23.32 | 37.38 | 15.91 | 30.74 | 0.075 |
| *P. aeruginosa* | *dfrB5* | *sul2* | Sulfonamide | 0.89 | 0.08 | 0.88 | 0.91 | 51.88 | 19.51 | 48.01 | 55.76 | ND |
| *E. cloacaea* | *dfrA1* | *sul1* | Sulfonamide | 0.37 | 0.23 | 0.32 | 0.42 | 10.68 | 7.19 | 9.25 | 12.10 | 1.24 |
| *E. cloacaea* | *dfrA14* | *sul2* | Sulfonamide | 0.30 | 0.11 | 0.28 | 0.32 | 14.69 | 7.62 | 13.17 | 16.20 | 1.62 |

**Table S4.** Co-occurrence of colistin (Gene A) and other antibiotic resistance genes in *Escherichia coli*. ND = not determined, SD = standard deviation, CI = confidence interval, BDPS = bidirectional probability score.

| **Gene A** | **Gene B** | **Gene B Resistance Type** | **Conditional Probability Mean** | **SD** | **CI (low)** | **CI (high)** | **Relative Risk** | **SD** | **CI (low)** | **CI (high)** | **BDPS** |
| --- | --- | --- | --- | --- | --- | --- | --- | --- | --- | --- | --- |
| *mcr-1* | *lnuF* | Lincosamide | 0.77 | 0.11 | 0.75 | 0.79 | 95.24 | 33.33 | 88.63 | 101.85 | ND |
| *mcr-1* | *aadA22* | Aminoglycoside | 0.39 | 0.12 | 0.37 | 0.41 | 41.57 | 17.1 | 38.18 | 44.97 | 1.04 |
| *mcr-1* | *tet(X4)* | Tetracycline | 0.37 | 0.13 | 0.34 | 0.39 | 27.83 | 10.42 | 25.76 | 29.9 | 3.23 |
| *mcr-1* | *tet(M)* | Tetracycline | 0.37 | 0.15 | 0.34 | 0.4 | 29.21 | 12.89 | 26.65 | 31.77 | 2.22 |
| *mcr-1.1* | *lnuF* | Lincosamide | 0.78 | 0.12 | 0.76 | 0.81 | 111.96 | 47.65 | 102.5 | 121.41 | ND |
| *mcr-1.1* | *aadA22* | Aminoglycoside | 0.39 | 0.11 | 0.37 | 0.41 | 45.85 | 19.55 | 41.97 | 49.73 | 3.16 |
| *mcr-1.1* | *tet(X4)* | Tetracycline | 0.38 | 0.13 | 0.36 | 0.41 | 29.81 | 9.8 | 27.87 | 31.76 | 2.1 |
| *mcr-1.1* | *tet(M)* | Tetracycline | 0.35 | 0.14 | 0.33 | 0.38 | 29.88 | 13.05 | 27.29 | 32.47 | 1.02 |
| *mcr-3.1* | *catA2* | Chloramphenicol | 0.35 | 0.25 | 0.3 | 0.4 | 304.28 | 586.69 | 187.87 | 420.69 | ND |
| *mcr-3.1* | *sul3* | Sulfonamide | 0.07 | 0.05 | 0.06 | 0.08 | 78.87 | 97.82 | 59.46 | 98.28 | 0.15 |
| *mcr-3.1* | *cmlA1* | Sulfonamide | 0.07 | 0.05 | 0.06 | 0.08 | 68.88 | 80.78 | 52.85 | 84.91 | 0.16 |
| *mcr-3.1* | *bleO* | Bleomycin | 0.06 | 0.04 | 0.06 | 0.07 | 120.15 | 154.18 | 89.56 | 150.74 | 0.11 |

**Table S5.** Common patterns of resistance gene co-occurrence in Gram-positive bacterial pathogens*.* ND = not determined, SD = standard deviation, CI = confidence interval, BDPS = bidirectional probability score.

| **Organism** | **Gene A** | **Gene B Resistance Type** | **Gene B** | **Gene B Resistance Type** | **Conditional Probability** | **SD** | **CI (low)** | **CI (high)** | **Relative Risk** | **SD** | **CI (low)** | **CI (high)** | **BDPS** |
| --- | --- | --- | --- | --- | --- | --- | --- | --- | --- | --- | --- | --- | --- |
| *S. aureus* | *fexA* | Phenicol | *spw* | Aminoglycoside | 0.87 | 0.09 | 0.85 | 0.88 | 257.95 | 190.48 | 220.16 | 295.75 | ND |
| *S. aureus* | *fexA* | Phenicol | *gyrA-S84A* | Quinolone | 0.74 | 0.11 | 0.72 | 0.76 | 121.57 | 56.70 | 110.32 | 132.82 | 1.51 |
| *S. aureus* | *fexA* | Phenicol | *lnuB* | Lincosamide | 0.59 | 0.11 | 0.57 | 0.62 | 138.89 | 76.84 | 123.64 | 154.13 | 0.94 |
| *S. aureus* | *fexA* | Phenicol | *lsaE* | Lincosamide/Streptogramin | 0.57 | 0.11 | 0.55 | 0.59 | 106.53 | 47.31 | 97.14 | 115.92 | 0.99 |
| *S. aureus* | *mphC* | Macrolide | *ant(6)-Ia* | Streptomycin | 0.52 | 0.04 | 0.52 | 0.53 | 829.53 | 756.12 | 679.50 | 979.56 | 0.53 |
| *S. aureus* | *ermA* | Macrolide | *ant(9)-Ia* | Aminoglycoside | 0.96 | 0.01 | 0.96 | 0.96 | 906.36 | 565.14 | 794.22 | 1018.49 | 0.96 |
| *S. aureus* | *ermA* | Macrolide | *parC-E84G* | quinolone | 0.95 | 0.02 | 0.95 | 0.95 | 7.78 | 0.70 | 7.64 | 7.92 | 2.08 |
| *S. aureus* | *ermA* | Macrolide | *mecI* | Methicillin | 0.90 | 0.02 | 0.90 | 0.91 | 15.55 | 2.27 | 15.10 | 16.00 | 1.19 |
| *S. aureus* | *ermB* | Macrolide | *tet(S)* | Tetracycline | 0.95 | 0.04 | 0.94 | 0.95 | 63.62 | 16.61 | 60.32 | 66.92 | ND |
| *S. aureus* | *ermB* | Macrolide | *vanH-A* | Vancomycin | 0.94 | 0.04 | 0.93 | 0.95 | 61.55 | 14.70 | 58.63 | 64.47 | ND |
| *S. aureus* | *ermB* | Macrolide | *dfrE* | Trimethoprim | 0.94 | 0.06 | 0.93 | 0.95 | 59.06 | 16.09 | 55.86 | 62.25 | ND |
| *S. aureus* | *ermB* | Macrolide | *vanZ-A* | Vancomycin | 0.94 | 0.04 | 0.93 | 0.95 | 62.69 | 16.55 | 59.41 | 65.98 | ND |
| *S. aureus* | *ermB* | Macrolide | *catA* | Chloramphenicol | 0.94 | 0.04 | 0.93 | 0.95 | 66.34 | 19.11 | 62.55 | 70.13 | ND |
| *S. aureus* | *ermB* | Macrolide | *vanY-A* | Vancomycin | 0.94 | 0.06 | 0.93 | 0.95 | 60.75 | 15.09 | 57.75 | 63.74 | ND |
| *S. aureus* | *ermB* | Macrolide | *vanS-A* | Vancomycin | 0.93 | 0.04 | 0.92 | 0.94 | 62.32 | 16.44 | 59.06 | 65.58 | ND |
| *S. aureus* | *ermB* | Macrolide | *vanR-A* | Vancomycin | 0.93 | 0.11 | 0.91 | 0.95 | 61.47 | 16.56 | 58.18 | 64.76 | ND |
| *S. aureus* | *ermB* | Macrolide | *vanX-A* | Vancomycin | 0.92 | 0.10 | 0.90 | 0.94 | 60.24 | 16.98 | 56.87 | 63.61 | ND |
| *S. aureus* | *ermB* | Macrolide | *vanA* | Vancomycin | 0.91 | 0.11 | 0.89 | 0.93 | 59.62 | 16.46 | 56.35 | 62.88 | ND |
| *S. aureus* | *mecA* | β-lactam | *gyrA-S84L* | Quinolone | 0.97 | 0.01 | 0.96 | 0.97 | 1.90 | 0.07 | 1.89 | 1.92 | 1.77 |
| *S. aureus* | *mecI* | β-lactam | *rpoB-S486L* | Rifamycin | 0.92 | 0.07 | 0.91 | 0.93 | 5.50 | 0.52 | 5.40 | 5.60 | ND |
| *S. aureus* | *mecI* | β-lactam | *parC-E84G* | Quinolone | 0.85 | 0.03 | 0.85 | 0.86 | 8.76 | 0.75 | 8.61 | 8.91 | 1.75 |
| *E. faecalis* | *fexA* | Phenicol | *optrA* | Oxazolidinone | 0.63 | 0.07 | 0.61 | 0.64 | 55.10 | 31.33 | 48.88 | 61.32 | 0.89 |
| *E. faecalis* | *fexA* | Phenicol | *ant(9)-Ia* | Aminoglycoside/Spectinomycin | 0.62 | 0.07 | 0.61 | 0.64 | 30.00 | 9.76 | 28.07 | 31.94 | 1.19 |
| *E. faecalis* | *fexA* | Phenicol | *gyrA-E87G* | Quinolone | 0.20 | 0.08 | 0.19 | 0.22 | 5.06 | 2.22 | 4.62 | 5.50 | 1.77 |
| *E. faecalis* | *ermB* | Macrolide | *ant(6)-Ia* | Streptomycin | 0.95 | 0.03 | 0.95 | 0.96 | 20.33 | 4.96 | 19.35 | 21.32 | 1.36 |
| *E. faecalis* | *ermB* | Macrolide | *aph(3’)-IIIa* | Amikacin/Kanamycin | 0.93 | 0.03 | 0.93 | 0.94 | 14.39 | 2.97 | 13.80 | 14.98 | 1.59 |
| *E. faecalis* | *ermB* | Macrolide | *lnuB* | Lincosamide | 0.93 | 0.03 | 0.93 | 0.94 | 9.64 | 1.34 | 9.37 | 9.90 | 2.53 |
| *E. faecalis* | *ermB* | Macrolide | *sat4* | Streptothricin | 0.93 | 0.03 | 0.92 | 0.94 | 12.98 | 2.54 | 12.48 | 13.49 | 1.72 |
| *E. faecalis* | *ermB* | Macrolide | *tet(L)* | Tetracycline | 0.69 | 0.07 | 0.68 | 0.71 | 6.11 | 0.92 | 5.93 | 6.30 | 2.55 |
| *E. faecium* | *fexA* | Phenicol | *lnuG* | Lincosamide | 0.20 | 0.15 | 0.17 | 0.23 | 10.30 | 8.64 | 8.58 | 12.01 | 1.89 |
| *E. faecium* | *ermA* | Macrolide | *an(t9)-Ia* | Aminoglycoside/Spectinomycin | 0.75 | 0.12 | 0.72 | 0.77 | 205.41 | 59.47 | 193.62 | 217.21 | 0.86 |
| *E. faecium* | *ermB* | Macrolide | *aph(3’)-IIIa* | Amikacin/Kanamycin | 0.72 | 0.06 | 0.70 | 0.73 | 40.07 | 12.19 | 37.66 | 42.49 | 1.36 |
| *E. faecium* | *ermB* | Macrolide | *sat4* | Streptothricin | 0.50 | 0.04 | 0.49 | 0.51 | 18.31 | 5.06 | 17.31 | 19.31 | 1.81 |

**Table S6.** Co-occurrence of vancomycin (Gene A) and other resistance genes in Gram-positive bacterial pathogens. ND = not determined, SD = standard deviation, CI = confidence interval, BDPS = bidirectional probability score.

| **Organism** | **Gene A** | **Gene B** | **Gene B Resistance Type** | **Conditional Probability** | **SD** | **CI (low)** | **CI (high)** | **Relative Risk** | **SD** | **CI (low)** | **CI (high)** | **BDPS** |
| --- | --- | --- | --- | --- | --- | --- | --- | --- | --- | --- | --- | --- |
| *S. aureus* | *vanA* | *vgaE* | Lincosamide | 0.15 | 0.10 | 0.13 | 0.17 | 37.87 | 36.49 | 30.63 | 45.11 | 2.38 |
| *S. aureus* | *vanA* | *gyrA-E88A* | Quinolone | 0.12 | 0.07 | 0.11 | 0.14 | 32.19 | 23.26 | 27.58 | 36.81 | 1.07 |
| *S. aureus* | *vanA* | *aph(3’)-IIIa* | Aminoglycoside | 0.04 | 0.02 | 0.04 | 0.04 | 34.34 | 21.26 | 30.12 | 38.56 | 0.05 |
| *S. aureus* | *vanH-A* | *vgaE* | Lincosamide | 0.15 | 0.12 | 0.13 | 0.18 | 47.12 | 75.50 | 32.14 | 62.10 | 2.18 |
| *S. aureus* | *vanH-A* | *gyrA-E88A* | Quinolone | 0.13 | 0.10 | 0.11 | 0.15 | 36.52 | 44.48 | 27.70 | 45.35 | 1.21 |
| *S. aureus* | *vanH-A* | *parC-E84K* | Quinolone | 0.11 | 0.05 | 0.10 | 0.12 | 67.04 | 24.61 | 62.16 | 71.92 | 0.16 |
| *S. aureus* | *vanH-A* | *catA* | Chloramphenicol | 0.05 | 0.04 | 0.04 | 0.06 | 14.27 | 15.74 | 11.15 | 17.39 | 0.83 |
| *S. aureus* | *vanH-A* | *aph(3’)-IIIa* | Aminoglycoside | 0.04 | 0.01 | 0.04 | 0.04 | 34.29 | 21.15 | 30.09 | 38.48 | 0.05 |
| *S. aureus* | *vanR-A* | *vgaE* | Lincosamide | 0.16 | 0.13 | 0.13 | 0.19 | 41.54 | 54.66 | 30.70 | 52.38 | 2.42 |
| *S. aureus* | *vanR-A* | *gyrA-E88A* | Quinolone | 0.13 | 0.08 | 0.11 | 0.15 | 35.95 | 27.54 | 30.48 | 41.41 | 1.18 |
| *S. aureus* | *vanR-A* | *parC-E84K* | Quinolone | 0.11 | 0.04 | 0.10 | 0.12 | 70.12 | 30.64 | 64.04 | 76.20 | 0.16 |
| *S. aureus* | *vanR-A* | *dfrE* | Trimethoprim | 0.05 | 0.10 | 0.03 | 0.07 | 17.21 | 60.28 | 5.25 | 29.17 | 1.91 |
| *S. aureus* | *vanR-A* | *aph(3’)-IIIa* | Aminoglycoside | 0.04 | 0.02 | 0.03 | 0.04 | 29.61 | 15.82 | 26.48 | 32.75 | 0.05 |
| *S. aureus* | *vanS-A* | *vgaE* | Lincosamide | 0.18 | 0.13 | 0.15 | 0.20 | 49.62 | 66.13 | 36.50 | 62.74 | 2.84 |
| *S. aureus* | *vanS-A* | *gyrA-E88A* | Quinolone | 0.13 | 0.08 | 0.12 | 0.15 | 35.76 | 35.53 | 28.71 | 42.81 | 1.21 |
| *S. aureus* | *vanS-A* | *parC-E84K* | Quinolone | 0.11 | 0.04 | 0.10 | 0.12 | 69.37 | 28.58 | 63.70 | 75.04 | 0.16 |
| *S. aureus* | *vanS-A* | *glpT-L27F* | Fosfomycin | 0.05 | 0.05 | 0.04 | 0.06 | 18.04 | 34.38 | 11.22 | 24.87 | 1.06 |
| *S. aureus* | *vanS-A* | *dfrE* | Trimethoprim | 0.05 | 0.09 | 0.03 | 0.07 | 17.50 | 53.70 | 6.84 | 28.15 | ND |
| *S. aureus* | *vanS-A* | *aph(3’)-IIIa* | Aminoglycoside | 0.04 | 0.02 | 0.03 | 0.04 | 34.30 | 22.67 | 29.81 | 38.80 | 0.05 |
| *S. aureus* | *vanX-A* | *vgaE* | Lincosamide | 0.16 | 0.11 | 0.14 | 0.18 | 40.16 | 38.21 | 32.57 | 47.74 | 2.30 |
| *S. aureus* | *vanX-A* | *gyrA-E88A* | Quinolone | 0.13 | 0.08 | 0.11 | 0.14 | 33.31 | 33.64 | 26.64 | 39.99 | 1.13 |
| *S. aureus* | *vanX-A* | *parC-E84K* | Quinolone | 0.11 | 0.04 | 0.10 | 0.11 | 72.39 | 57.09 | 61.06 | 83.72 | 0.16 |
| *S. aureus* | *vanX-A* | *glpT-L27F* | Fosfomycin | 0.06 | 0.05 | 0.05 | 0.07 | 18.01 | 27.33 | 12.58 | 23.43 | 1.08 |
| *S. aureus* | *vanX-A* | *aph(3’)-IIIa* | Aminoglycoside | 0.04 | 0.02 | 0.04 | 0.04 | 39.88 | 51.85 | 29.59 | 50.17 | 0.05 |
| *S. aureus* | *vanY-A* | *vgaE* | Lincosamide | 0.15 | 0.11 | 0.13 | 0.18 | 46.93 | 81.98 | 30.67 | 63.20 | 2.58 |
| *S. aureus* | *vanY-A* | *gyrA-E88A* | Quinolone | 0.13 | 0.08 | 0.11 | 0.15 | 35.81 | 32.77 | 29.31 | 42.31 | 1.18 |
| *S. aureus* | *vanY-A* | *parC-E84K* | Quinolone | 0.11 | 0.04 | 0.10 | 0.11 | 64.81 | 25.49 | 59.75 | 69.87 | 0.16 |
| *S. aureus* | *vanY-A* | *glpT-L27F* | Fosfomycin | 0.07 | 0.06 | 0.05 | 0.08 | 24.68 | 66.96 | 11.39 | 37.96 | 1.13 |
| *S. aureus* | *vanY-A* | *catA* | Chloramphenicol | 0.05 | 0.03 | 0.04 | 0.05 | 16.99 | 54.88 | 6.10 | 27.88 | 0.82 |
| *S. aureus* | *vanY-A* | *aph(3’)-IIIa* | Aminoglycoside | 0.04 | 0.02 | 0.04 | 0.04 | 33.29 | 24.15 | 28.50 | 38.09 | 0.05 |
| *S. aureus* | *vanZ-A* | *vgaE* | Lincosamide | 0.16 | 0.12 | 0.14 | 0.19 | 45.87 | 82.83 | 29.44 | 62.30 | 2.80 |
| *S. aureus* | *vanZ-A* | *parC-E84K* | Quinolone | 0.11 | 0.04 | 0.10 | 0.12 | 69.79 | 29.83 | 63.87 | 75.71 | 0.16 |
| *S. aureus* | *vanZ-A* | *catA* | Chloramphenicol | 0.05 | 0.04 | 0.04 | 0.05 | 11.98 | 11.86 | 9.63 | 14.33 | 0.64 |
| *S. aureus* | *vanZ-A* | *gyrA-S84V* | Quinolone | 0.04 | 0.10 | 0.02 | 0.06 | 14.55 | 54.57 | 3.72 | 25.38 | 1.40 |
| *S. aureus* | *vanZ-A* | *aph(3’)-IIIa* | Aminoglycoside | 0.04 | 0.02 | 0.04 | 0.04 | 34.53 | 26.00 | 29.37 | 39.69 | 0.05 |
| *E. faecium* | *vanB* | *mefH* | Macrolide | 0.29 | 0.07 | 0.28 | 0.31 | 5.47 | 1.36 | 5.20 | 5.74 | 1.15 |
| *E. faecium* | *vanB* | *gyrA-S83I* | Quinolone | 0.26 | 0.05 | 0.25 | 0.27 | 6.99 | 2.21 | 6.55 | 7.43 | 0.53 |
| *E. faecium* | *vanB* | *dfrG* | Trimethoprim | 0.23 | 0.04 | 0.23 | 0.24 | 8.73 | 2.82 | 8.17 | 9.29 | 0.35 |
| *E. faecium* | *vanH-B* | *mefH* | Macrolide | 0.29 | 0.06 | 0.28 | 0.30 | 5.44 | 1.24 | 5.20 | 5.69 | 1.15 |
| *E. faecium* | *vanH-B* | *gyrA-S83I* | Quinolone | 0.27 | 0.05 | 0.25 | 0.28 | 6.98 | 2.12 | 6.56 | 7.40 | 0.53 |
| *E. faecium* | *vanH-B* | *dfrG* | Trimethoprim | 0.23 | 0.04 | 0.23 | 0.24 | 8.55 | 2.68 | 8.02 | 9.08 | 0.35 |
| *E. faecium* | *vanR-A* | *sat4* | Streptothricin | 0.47 | 0.05 | 0.46 | 0.48 | 26.64 | 6.12 | 25.43 | 27.86 | 1.36 |
| *E. faecium* | *vanR-A* | *ant6)-Ia* | Aminoglycoside | 0.38 | 0.06 | 0.37 | 0.39 | 17.98 | 5.55 | 16.88 | 19.08 | 1.64 |
| *E. faecium* | *vanR-A* | *aph(3’)-IIIa* | Aminoglycoside | 0.30 | 0.05 | 0.29 | 0.31 | 15.34 | 4.45 | 14.46 | 16.22 | 0.99 |
| *E. faecium* | *vanR-A* | *23S-G2576T* | Oxazolidinone | 0.28 | 0.15 | 0.25 | 0.31 | 11.93 | 6.85 | 10.57 | 13.29 | 1.63 |
| *E. faecium* | *vanR-A* | *ermB* | Macrolide | 0.26 | 0.05 | 0.25 | 0.27 | 14.68 | 4.06 | 13.88 | 15.49 | 0.73 |
| *E. faecium* | *vanS-A* | *sat4* | Streptothricin | 0.49 | 0.04 | 0.48 | 0.50 | 31.85 | 10.04 | 29.86 | 33.85 | 1.24 |
| *E. faecium* | *vanS-A* | *ant6-Ia* | Aminoglycoside | 0.42 | 0.08 | 0.41 | 0.44 | 23.27 | 9.45 | 21.40 | 25.14 | 1.55 |
| *E. faecium* | *vanS-A* | *aph(3’)-IIIa* | Aminoglycoside | 0.33 | 0.05 | 0.32 | 0.34 | 20.40 | 9.31 | 18.55 | 22.25 | 0.93 |
| *E. faecium* | *vanS-A* | *ermB* | Macrolide | 0.28 | 0.04 | 0.27 | 0.29 | 18.58 | 6.74 | 17.24 | 19.92 | 0.68 |
| *E. faecium* | *vanS-A* | *23S-G2576T* | Oxazolidinone | 0.27 | 0.15 | 0.24 | 0.30 | 13.11 | 8.34 | 11.46 | 14.76 | 1.47 |
| *E. faecium* | *vanS-B* | *mefH* | Macrolide | 0.30 | 0.06 | 0.28 | 0.31 | 5.42 | 1.30 | 5.17 | 5.68 | 1.16 |
| *E. faecium* | *vanS-B* | *gyrA-S83I* | Quinolone | 0.26 | 0.05 | 0.25 | 0.27 | 6.82 | 2.07 | 6.41 | 7.23 | 0.53 |
| *E. faecium* | *vanS-B* | *dfrG* | Trimethoprim | 0.23 | 0.04 | 0.23 | 0.24 | 8.40 | 2.40 | 7.92 | 8.88 | 0.35 |
| *E. faecium* | *vanW-B* | *dfrG* | Trimethoprim | 0.17 | 0.03 | 0.17 | 0.18 | 6.96 | 2.07 | 6.55 | 7.37 | 0.28 |
| *E. faecium* | *vanX-B* | *mefH* | Macrolide | 0.29 | 0.07 | 0.28 | 0.31 | 5.45 | 1.29 | 5.19 | 5.70 | 1.15 |
| *E. faecium* | *vanX-B* | *gyrA-S83I* | Quinolone | 0.26 | 0.05 | 0.25 | 0.28 | 7.05 | 2.20 | 6.61 | 7.48 | 0.53 |
| *E. faecium* | *vanX-B* | *dfrG* | Trimethoprim | 0.23 | 0.04 | 0.22 | 0.24 | 8.74 | 2.70 | 8.20 | 9.27 | 0.35 |
| *E. faecium* | *vanY-B* | *mefH* | Macrolide | 0.30 | 0.06 | 0.28 | 0.31 | 5.46 | 1.26 | 5.21 | 5.71 | 1.15 |
| *E. faecium* | *vanY-B* | *gyrA-S83I* | Quinolone | 0.27 | 0.06 | 0.25 | 0.28 | 7.01 | 2.11 | 6.60 | 7.43 | 0.53 |
| *E. faecium* | *vanY-B* | *dfrG* | Trimethoprim | 0.23 | 0.04 | 0.23 | 0.24 | 8.56 | 2.47 | 8.07 | 9.05 | 0.35 |
| *E. faecalis* | *vanB* | *fexB* | Phenicol | 0.10 | 0.05 | 0.09 | 0.11 | 10.69 | 7.85 | 9.14 | 12.25 | 0.98 |
| *E. faecalis* | *vanB* | *tet(O-W-32-O)* | Tetracycline | 0.09 | 0.05 | 0.08 | 0.10 | 9.10 | 7.07 | 7.69 | 10.50 | 1.50 |
| *E. faecalis* | *vanD* | *fexB* | Phenicol | 0.09 | 0.04 | 0.08 | 0.10 | 13.51 | 14.65 | 10.60 | 16.41 | 0.76 |
| *E. faecalis* | *vanD* | *tet(O-W-32-O)* | Tetracycline | 0.09 | 0.04 | 0.08 | 0.10 | 11.80 | 11.09 | 9.60 | 14.00 | 1.10 |
| *E. faecalis* | *vanD* | *gyrA-S83I* | Quinolone | 0.04 | 0.02 | 0.04 | 0.04 | 7.77 | 1.84 | 7.41 | 8.14 | 0.08 |
| *E. faecalis* | *vanH-B* | *fexB* | Phenicol | 0.09 | 0.04 | 0.08 | 0.10 | 9.69 | 7.26 | 8.25 | 11.13 | 0.88 |
| *E. faecalis* | *vanH-B* | *tet(O-W-32-O)* | Tetracycline | 0.08 | 0.04 | 0.08 | 0.09 | 8.77 | 6.89 | 7.40 | 10.14 | 1.43 |
| *E. faecalis* | *vanH-D* | *fexB* | Phenicol | 0.08 | 0.03 | 0.08 | 0.09 | 11.70 | 10.11 | 9.70 | 13.71 | 0.71 |
| *E. faecalis* | *vanH-D* | *tet(O-W-32-O)* | Tetracycline | 0.08 | 0.03 | 0.07 | 0.09 | 10.30 | 7.84 | 8.75 | 11.86 | 1.06 |
| *E. faecalis* | *vanH-D* | *gyrA-S83I* | Quinolone | 0.04 | 0.02 | 0.04 | 0.04 | 7.84 | 2.28 | 7.39 | 8.29 | 0.08 |
| *E. faecalis* | *vanR-B* | *fexB* | Phenicol | 0.10 | 0.04 | 0.09 | 0.11 | 10.52 | 7.38 | 9.05 | 11.98 | 1.00 |
| *E. faecalis* | *vanR-B* | *tet(O-W-32-O)* | Tetracycline | 0.09 | 0.05 | 0.08 | 0.10 | 8.53 | 6.38 | 7.26 | 9.79 | 1.42 |
| *E. faecalis* | *vanR-D* | *tet(O-W-32-O)* | Tetracycline | 0.08 | 0.04 | 0.08 | 0.09 | 11.84 | 14.01 | 9.06 | 14.62 | 1.09 |
| *E. faecalis* | *vanR-D* | *fexB* | Phenicol | 0.08 | 0.04 | 0.07 | 0.09 | 10.83 | 7.76 | 9.29 | 12.37 | 0.77 |
| *E. faecalis* | *vanR-D* | *gyrA-S83I* | Quinolone | 0.04 | 0.02 | 0.04 | 0.04 | 7.81 | 1.97 | 7.42 | 8.21 | 0.08 |
| *E. faecalis* | *vanS-B* | *fexB* | Phenicol | 0.10 | 0.04 | 0.09 | 0.11 | 10.64 | 8.53 | 8.95 | 12.33 | 0.96 |
| *E. faecalis* | *vanS-B* | *tet(O-W-32-O)* | Tetracycline | 0.09 | 0.05 | 0.08 | 0.10 | 8.66 | 7.06 | 7.26 | 10.06 | 1.44 |
| *E. faecalis* | *vanS-D* | *tet(O-W-32-O)* | Tetracycline | 0.08 | 0.04 | 0.07 | 0.08 | 10.42 | 8.42 | 8.75 | 12.09 | 0.91 |
| *E. faecalis* | *vanS-D* | *fexB* | Phenicol | 0.07 | 0.04 | 0.06 | 0.08 | 8.49 | 5.77 | 7.34 | 9.63 | 0.85 |
| *E. faecalis* | *vanS-D* | *gyrA-S83I* | Quinolone | 0.04 | 0.02 | 0.04 | 0.04 | 8.63 | 2.23 | 8.18 | 9.07 | 0.07 |
| *E. faecalis* | *vanW-B* | *fexB* | Phenicol | 0.11 | 0.04 | 0.10 | 0.12 | 10.70 | 6.40 | 9.42 | 11.97 | 1.00 |
| *E. faecalis* | *vanW-B* | *tet(O-W-32-O)* | Tetracycline | 0.08 | 0.04 | 0.07 | 0.09 | 7.64 | 5.32 | 6.58 | 8.69 | 1.63 |
| *E. faecalis* | *vanX-B* | *fexB* | Phenicol | 0.12 | 0.05 | 0.11 | 0.13 | 11.14 | 6.99 | 9.75 | 12.53 | 1.01 |
| *E. faecalis* | *vanX-B* | *tet(O-W-32-O)* | Tetracycline | 0.10 | 0.05 | 0.09 | 0.11 | 8.28 | 5.36 | 7.22 | 9.35 | 1.58 |
| *E. faecalis* | *vanX-D* | *tet(O-W-32-O)* | Tetracycline | 0.05 | 0.03 | 0.05 | 0.06 | 7.42 | 8.96 | 5.64 | 9.20 | 1.01 |
| *E. faecalis* | *vanY-B* | *fexB* | Phenicol | 0.10 | 0.04 | 0.09 | 0.11 | 10.61 | 7.59 | 9.11 | 12.12 | 0.95 |
| *E. faecalis* | *vanY-B* | *tet(O-W-32-O)* | Tetracycline | 0.09 | 0.04 | 0.08 | 0.10 | 8.62 | 5.52 | 7.52 | 9.71 | 1.49 |

**Table S7.** Co-occurrence of heavy metal and antibiotic resistance genes in Gram-negative bacterial pathogens and *S. aureus*. Due to the size of this table, it is available as a separate downloadable file from https://github.com/ERBringHorvath/regain_cl/tree/main/probability_tables.

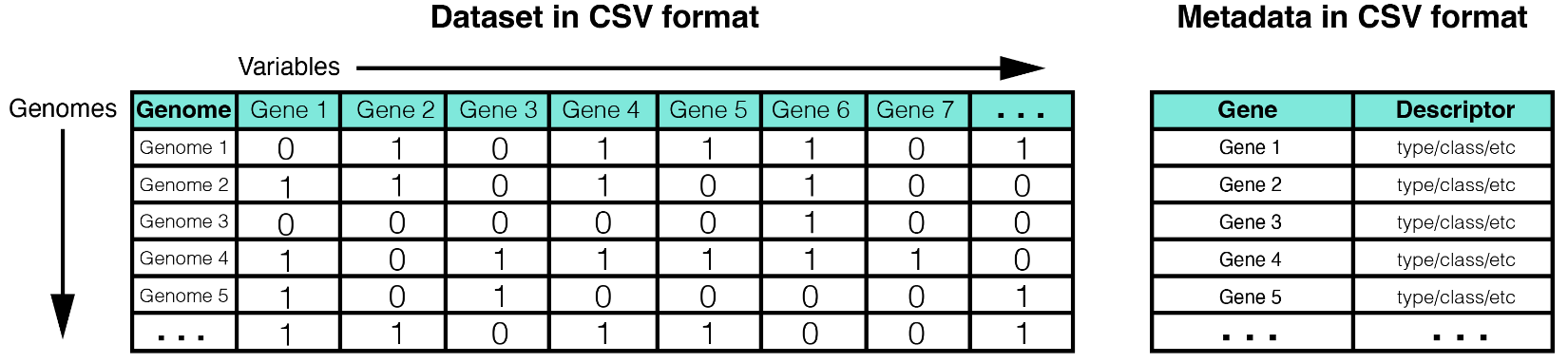

**Figure S1**. Required format of externally prepared data. The data matrix must be in presence/absence format, where ‘1’ = present and ‘0’ = absent. Rows and columns must be labeled by genome and gene, respectively. Each variable must have two states (i.e., both '1' and '0'). Including variables that have only one state, such as a gene that is present in every genome will result in the failure of the Bayesian network analysis. Variables may only contain alphanumeric characters and underscores (Do NOT include quotation marks, periods, forward or reverse slash marks, hyphens, parentheses). Addition of these characters will result in the variable being excluded from the Bayesian network analysis. Metadata should have two columns, one containing the names of your variables and the second containing variable description data. We suggest using gene family or class as a descriptor. All data files must be in CSV format. If you are working in Excel, data tables can be exported to CSV via: File > Save As, then select 'Comma Separated Values (.csv)' from the drop-down menu.

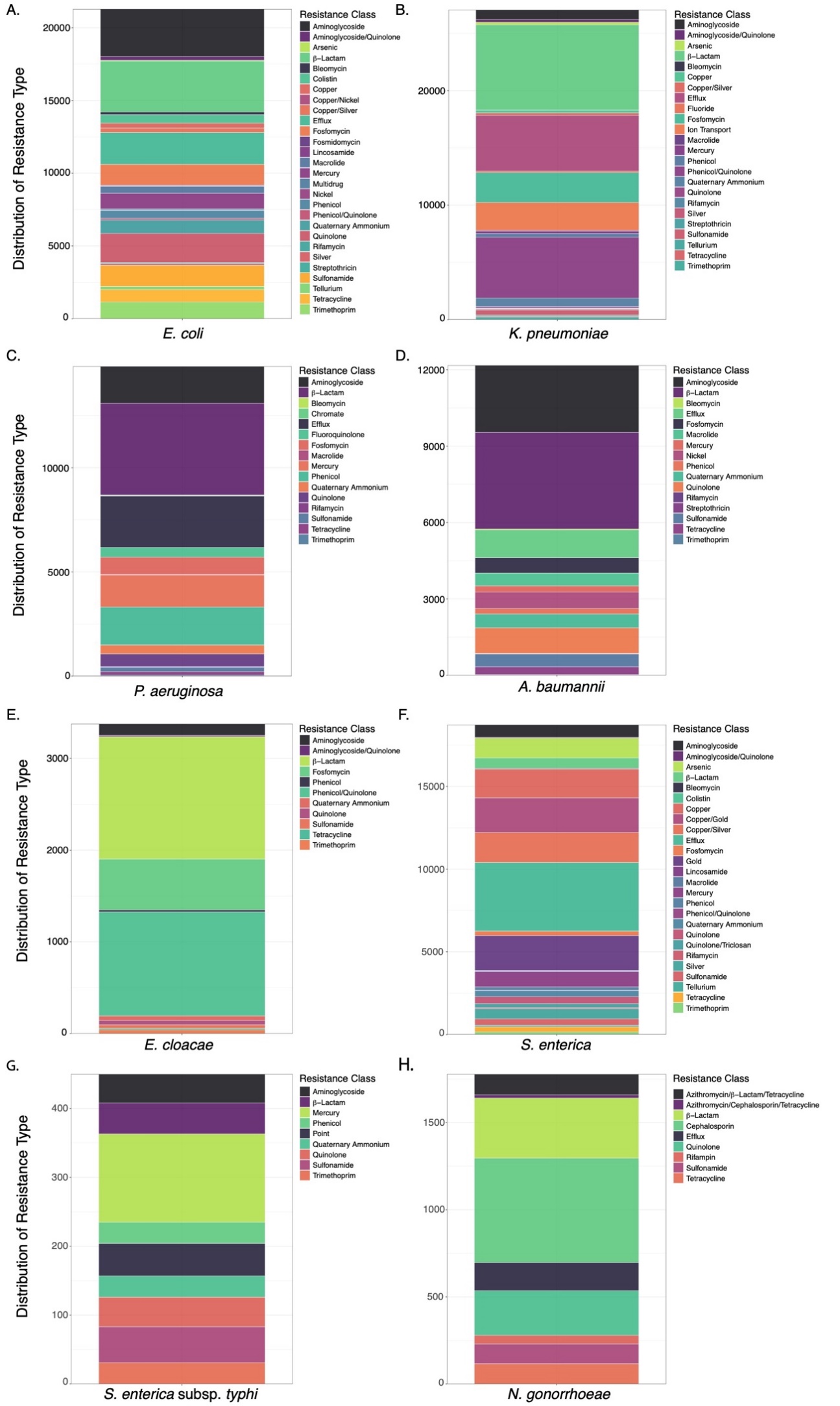

**Figure S2.** Distribution of resistance gene classes across each Gram-negative bacterial genomic population. **A−H.** Resistance gene classes identified in each genomic population. **A.** *Escherichia coli*. **B.** *Klebsiella pneumoniae*. **C.** *Pseudomonas aeruginosa*. **D.** *Acinetobacter baumannii*. **E.** *Enterobacter cloacae*. **F.** *Salmonella enterica.* **G.** *Salmonella enterica* subsp. *typhi*. **H.** *Neisseria gonorrhoeae*.

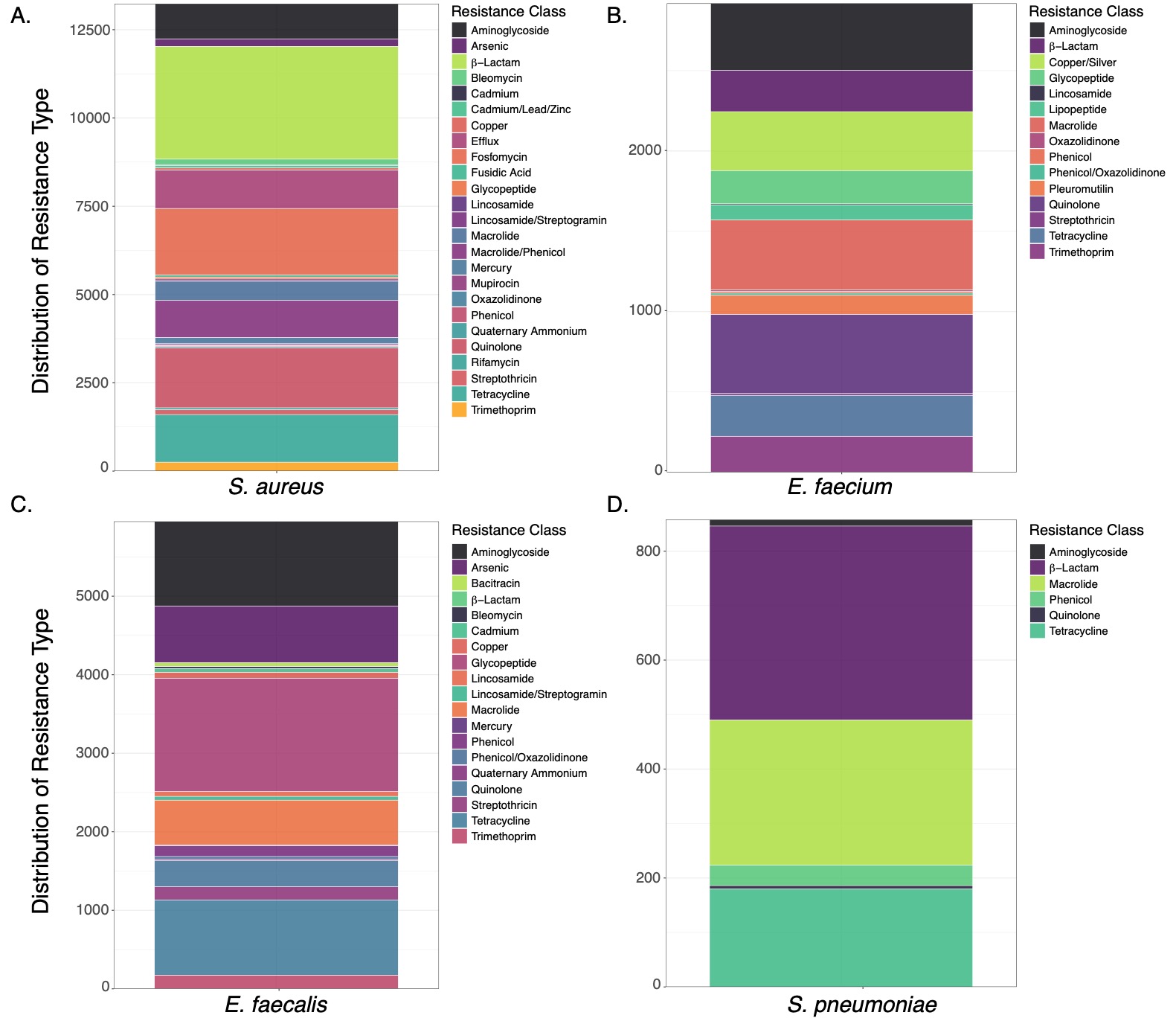

**Figure S3.** Distribution of resistance gene classes across each Gram-positive bacterial genomic population. **A−D.** Resistance gene classes identified in each genomic population. **A.** *Staphylococcus aureus*. **B.** *Enterococcus faecium*. **C.** *Enterococcus faecalis*. **D.** *Streptococcus pneumoniae*.

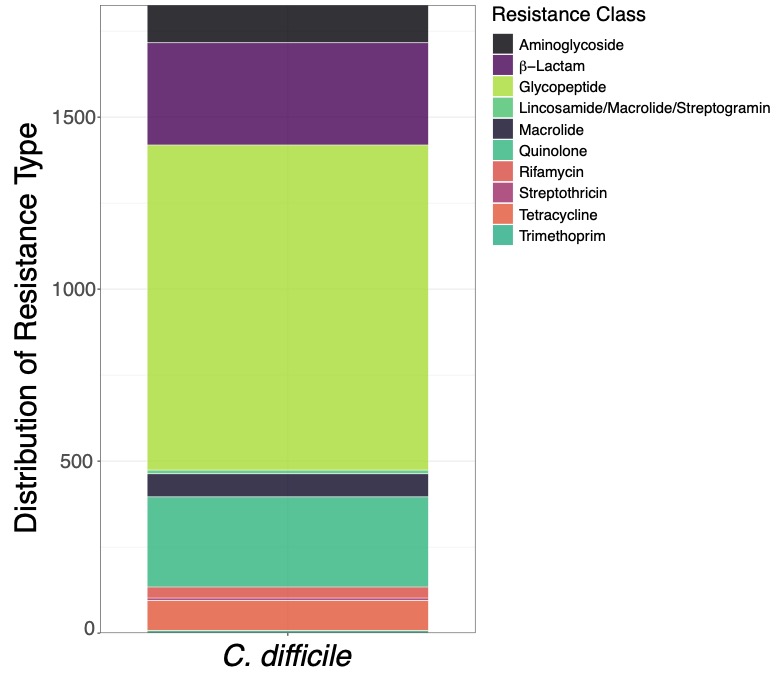

**Figure S4.** Resistance gene classes identified in the *Clostridium difficile* genomic population.
